# Investigating robust associations between functional connectivity based on graph theory and general intelligence

**DOI:** 10.1101/2023.07.18.549314

**Authors:** Dorothea Metzen, Christina Stammen, Christoph Fraenz, Caroline Schlüter, Wendy Johnson, Onur Güntürkün, Colin G. DeYoung, Erhan Genç

**Affiliations:** Biopsychology, Institute for Cognitive Neuroscience, Faculty of Psychology, Ruhr University Bochum, 44801 Bochum, Germany; Department of Psychology and Neuroscience, Leibniz Research Centre for Working Environment and Human Factors (IfADo), 44139 Dortmund, Germany; Department of Psychology, University of Edinburgh, Edinburgh, EH8 9JZ, UK; Department of Psychology, University of Minnesota, Minneapolis, MN 55455, USA

**Author notes:** These authors contributed equally (Dorothea Metzen, Christina Stammen).

**Keywords:** resting-state fMRI, general intelligence, multi-center study, graph theory, functional network connectivity

## Abstract

Previous research investigating relations between general intelligence and graph-theoretical properties of the brain’s intrinsic functional network has yielded contradictory results. A promising approach to tackle such mixed findings is multi-center analysis. For this study, we analyzed data from four independent data sets (total N > 2000) to identify robust associations amongst samples between *g* factor scores and global as well as node-specific graph metrics. On the global level, *g* showed no significant associations with global efficiency in any sample, but significant positive associations with global clustering coefficient and small-world propensity in two samples. On the node-specific level, elastic-net regressions for nodal efficiency and local clustering yielded no brain areas that exhibited consistent associations amongst data sets. Using the areas identified via elastic-net regression in one sample to predict *g* in other samples was not successful for nodal efficiency and only led to significant predictions between two data sets for local clustering. Thus, using conventional graph theoretical measures based on resting-state imaging did not result in replicable associations between functional connectivity and general intelligence.

## 1. Introduction

Any system composed of interrelated elements can be modeled as a network ^[1]^. One of the most challenging and complex systems found in nature is the human brain ^[2, 3]^. It is not possible to study the whole brain’s connectivity at a cellular level using current in-vivo neuroimaging methods ^[4]^. Nevertheless, over the past four decades, structural and functional neuroimaging studies have generated an enormous amount of knowledge about the human brain at a macroscopic scale ^[5]^. For example, it has been observed that human brain networks are organized in a ‘small-world’ manner. That is, a high level of local clustering, which enables segregated functionally coherent units of local information processing, co-exists with robust long-distance connections, ensuring integration of information across different regions ^[1, 6–8]^. As no two individual’s brains are completely alike ^[3]^, many studies addressed the question whether interindividual differences in structural or functional brain properties are linked to interindividual differences in human behavior. Within the field of psychology, examining interindividual differences in the form of general cognitive performance (i.e. intelligence) exhibits the longest research tradition ^[9]^.

Intelligence can be defined as the “[…] ability to understand complex ideas, to adapt effectively to the environment, to learn from experience, to engage in various forms of reasoning, to overcome obstacles by taking thought” ^[10]^. Spearman ^[11]^ observed that individuals who perform above average on one cognitive task also tend to do well on other cognitive tasks. Based on this observation, he identified the existence of a ‘general factor’ of intelligence that he termed ‘*g’*. Current hierarchically organized models still place *g* at the top of the hierarchy with more specific broadly organized cognitive abilities underneath that are then further subdivided at lower levels ^[12]^. Interindividual differences in cognitive performance are not only relatively stable across tasks but also persist throughout the lifespan ^[13]^. Moreover, intelligence test scores predict many aspects of life and health such as job performance and longevity ^[10, 14–17]^. Given the considerable impact that *g* seems to have on life outcomes, the question “Where in the brain is intelligence?” ^[18]^ has always been of special interest.

Many neuroscientific studies attempted to answer this question by employing various imaging techniques to analyze differences in the neural properties of brain regions and relate them to intelligence. Combined evidence from such studies led three independent meta-analyses ^[18–20]^ to conclude that networks of brain areas widely distributed across the brain are associated with intelligence. Jung and Haier ^[18]^ suggested that intelligence-related areas comprise an interconnected, widespread network (instead of working in isolation from each other) and proposed the Parieto-Frontal Integration Theory of intelligence (P-FIT). The conception of P-FIT represents an important milestone in the endeavor of identifying the neural basis of human intelligence. However, many years of neuroscientific intelligence research have passed since Jung and Haier first outlined their P-FIT network and new ideas were added to the original model such as the importance of subcortical structures ^[19]^.

More recent studies have directed attention towards analysis of intelligence-related neural properties in terms of network organization ^[2, 21–23]^. One branch of network neuroscience ^[22]^ is focused on functional connectivity within the human brain. This approach emerges from the idea that resting-state functional connectivity of anatomically separated brain regions, quantified via functional magnetic resonance imaging (fMRI), reflects the brain’s fundamental architecture of functional networks underlying task-related activities and behavior ^[5, 24–26]^. Functional connectivity among brain regions can be inferred from temporal correlations between spontaneous low-frequency fluctuations in the blood oxygenation level-dependent (BOLD) signals ^[27, 28]^. Given that functional connectivity appears to be unique to individuals, almost comparable to fingerprints in reliability and robustness ^[29]^, studies began to focus on intelligence-related differences in the brain’s intrinsic functional organization by applying graph theory.

Graph theory, a branch of mathematics concerned with representing and characterizing complex networks, provides a powerful possibility to specify the brain’s topological organization ^[3, 30–32]^. One of the first studies employing it observed a negative association between intelligence quotient (IQ) scores and the characteristic path length of the resting-state brain network, indicating clear small-world organization ^[21]^. As its characteristic path length is inversely related to a network’s global efficiency ^[30]^, their results suggested that intelligence depends on how efficiently information is globally integrated among different brain regions. Van den Heuvel et al. ^[21]^ did not find a significant correlation between IQ scores and the global clustering coefficient, which measures the extent of local “cliquishness”, an indicator of locally segregated information processing ^[3, 7, 32]^. They also analyzed the topological properties of individual nodes (defined as single voxels) in relation to the rest of the brain. Voxels with significant correlations between their normalized path length and IQ scores were found in medial prefrontal gyrus, precuneus/posterior cingulate gyrus, bilateral inferior parietal regions, left superior temporal gyrus, and left inferior frontal gyrus. The observation that these voxels possessed more efficient (i.e. shorter) functional paths to other brain regions in more intelligent participants emphasized their relevance in the functional brain network. Later studies were able to provide further evidence of an association between intelligence and global efficiency by focusing their analyses on more specific aspects such as subnetworks, weak connections, or brain resilience ^[33–35]^.

In contrast, Pamplona et al. ^[36]^ did not observe a statistically significant relation between IQ scores and characteristic path length, global efficiency, or global clustering coefficient. On the level of individual network nodes, they reported significant correlations (uncorrected for multiple comparisons) between IQ scores and local efficiency values of bilateral pre-central regions as well as the left inferior occipital region. In line with the results of Pamplona et al. ^[36]^, Hilger et al. ^[37]^, who conceptualized their study similar to that of van den Heuvel et al. ^[21]^, did not observe any relation between IQ scores and global efficiency, but reported significant associations between IQ scores and nodal efficiency. They observed associations between IQ scores and nodal efficiency in right anterior insula and dorsal anterior cingulate cortex as well as lower nodal efficiency in the left temporo-parietal junction area. As noted by Kruschwitz et al. ^[38]^ among many others, the assumption that the brain’s global functional network efficiency is associated with general intelligence has been widely accepted, but the empirical foundation of this claim is rather weak. While aforementioned studies had rather small samples of 19 ^[21]^, 29 ^[36]^, and 54 participants ^[37]^, Kruschwitz et al. ^[38]^ used a large sample (n = 1096) and high quality data from the Human Connectome Project ^[39]^. They employed multiple network definitions and did not observe any significant associations between total cognition composite scores and characteristic path length, global efficiency, or global clustering coefficient ^[38]^.

Taken together, these results question the proposed association of general intelligence and the brain’s global functional network efficiency. Not only are there contradicting results on a global level, but results on a nodal level also do not overlap locally. As discussed by Kruschwitz et al. ^[38]^, methodological differences among studies limit their comparability and might contribute to differences in study observations. Our study utilized a multi-center approach, analyzing multiple, independent data sets to identify extents to which robust associations replicated across samples ^[40]^. In a recent study, we were able to find such associations between general intelligence and white matter microstructure by following this approach ^[41]^. Here, we examined associations between *g* and functional network properties in the same four independent data sets comprising more than 2000 healthy participants. All data sets were pre-processed and analyzed in the same manner to make them as comparable as possible. On the global level, we calculated associations between *g* and global efficiency, global clustering coefficient, and small-world propensity. On the nodal level, we used regularized elastic-net regressions to examine associations between *g* and nodal efficiency as well as local clustering coefficient of specific brain areas. To assess robustness of brain areas’ associations with general intelligence, we investigated whether the areas identified by the elastic-net analyses overlapped among data sets and tested the predictors identified in one sample in other samples. To test whether associations between graph metrics and *g* were affected by data reliability, we additionally investigated test-retest reliabilities of global and nodal metrics in two of the data sets.

## 2. Methods and Materials

Given that the data sets included in our study employed different behavioral measures and imaging data were obtained on different scanners, pooling into one big sample was not possible.

### 2.1. Participants

#### 2.1.1. RUB Sample

The RUB sample consisted of 557 participants aged 18-75 (mean age: 27.3 years, SD = 9.4 years, 274 women, 503 right-handers). Participants were either financially compensated or received course credit for their participation. All participants reported freedom from neurological and mental illnesses. Individuals were excluded from participation if they had insufficient German skills or reported familiarity with any of the tests used. The study was approved by the local ethics committee of the Faculty of Psychology at the Ruhr-University Bochum (vote 165). All participants gave written informed consent and were treated according to the Declaration of Helsinki.

#### 2.1.2. HCP Sample

Data were provided by the Human Connectome Project, WU-Minn Consortium (Principal Investigators: David Van Essen and Kamil Ugurbil; 1U54MH091657), funded by the 16 United States National Institutes of Health (NIH) Institutes and Centers supporting the NIH Blueprint for Neuroscience Research and by the McDonnell Center for Systems Neuroscience at Washington University. We used the “1200 Subjects Data Release” ^[39]^, which comprises behavioral and imaging data from 1206 young adults aged between 22 and 37. To calculate the *g* factor, all participants with missing data in any of the intelligence tests had to be excluded, which reduced the sample size to N = 1188 (mean age: 28.8 years, SD = 3.7 years, 641 females, 934 right-handers). HCP provides four resting-state functional magnetic resonance imaging (rsfMRI) scans from two different testing days (HCP_day_1 and HCP_day_2). However, not all rsfMRI data were available for all participants. Thus, the sample for HCP_day_1 was reduced to N = 1050 (mean age: 28.7 years, SD = 3.7 years, 564 females, 823 right-handers) and the sample for HCP_day_2 was reduced to N = 1011 (mean age: 28.7 years, SD = 3.7 years, 543 females, 794 right-handers). 1007 participants (mean age: 28.7 years, SD = 3.7 years, 539 females, 791 right-handers) completed all four rsfMRI scans (HCP_day_1_day_2). Participants reported no history of substance abuse or psychiatric, neurological, or cardiological diseases. All participants gave written informed consent ^[42]^.

#### 2.1.3. UMN Sample

The UMN sample consisted of 335 participants aged between 20 and 40 (mean age: 26.3 years, SD = 5.0 years, 164 females, all right-handed) and all participants were included in calculating the *g* factor. As rsfMRI data were not available for all participants, our sample was reduced to N = 274 (mean age: 26.2 years, SD = 4.9 years, 135 females, all right-handed). Exclusion criteria were neurological or psychiatric disorders, current drug abuse, and current use of psychotropic medication. The study protocol was approved by the University of Minnesota Institutional Review Board and all participants gave written informed consent.

#### 2.1.4. NKI Sample

The “Enhanced Nathan Kline Institute - Rockland Sample” data set ^[43]^ is part of the 1000 Functional Connectomes Project (http://fcon_1000.projects.nitrc.org) and was downloaded from its official website (http://fcon_1000.projects.nitrc.org/indi/enhanced/). We only included healthy participants, who reported freedom from mental illness. The *g* factor was calculated based on 417 participants aged between 6 and 85 (mean age: 43.5 years, SD = 23.5 years, 273 females, 326 right-handers). The three available rsfMRI measurements from NKI differ in temporal resolution (NKI_TR646, NKI_TR1400, and NKI_TR2500). After excluding all participants without rsfMRI data we ended up with N = 385 for NKI_TR645 aged 6-85 (mean age: 44.4 years, SD = 22.8 years, 252 females, 303 right-handers), N = 348 for NKI_TR1400 aged 6-83 (mean age: 44.5 years, SD = 22.6 years, 228 females, 270 right-handers) and N = 391 for NKI_TR2500 aged 6-85 (mean age: 44.4 years, SD = 22.8 years, 257 females, 308 right-handers). The study was approved by the Institutional Review Boards at the Nathan Kline Institute and Montclair State University. Participants and – if underaged – their legal guardians gave written informed consent.

### 2.2. Intelligence measurement

Detailed descriptions of all intelligence measures used to calculate *g* factors can be found in Stammen et al. ^[41]^. A short overview of all tests used is in the Supplementary Material (see “1. Intelligence measurement”).

### 2.3. Computation of the general intelligence factor (*g*)

As observed by Johnson et al. ^[44]^ and Johnson et al. ^[45]^, *g* factors derived from different test batteries are statistically equivalent, provided that included tests measure intelligence broadly enough. Since different test batteries were employed by our four samples, we decided to calculate *g* factors (one for each sample) to obtain intelligence measures comparable among data sets.

As described in Stammen et al. ^[41]^, we computed *g* factor scores based on each sample’s intelligence test scores (see Supplementary Material). After regressing age, sex, age*sex, age^2^, and age^2^*sex from the test scores, we conducted exploratory factor analyses based on the standardized residuals to develop hierarchical factor models for each sample. Following this, we performed confirmatory factor analyses and assessed model fit by the chi-square (Χ^2^) statistic as well as the fit indices Root Mean Square Error of Approximation (RMSEA), Standardized Root Mean Square Residual (SRMR), Comparative Fit Index (CFI), and Tucker-Lewis Index (TLI). From these models, we calculated regression-based *g*-factor scores for each participant, winsorizing outliers ^[46]^. The postulated confirmatory factor models for the four samples, the z-standardized factor loadings, and the covariances between individual subtests are shown in Figures 1-4 in Stammen et al. ^[41]^. The evaluation of model fit yielded good (samples RUB and HCP) to excellent (samples UMN and NKI) fit (see Table 2 in Stammen et al. ^[41]^).

**Figure 1.**
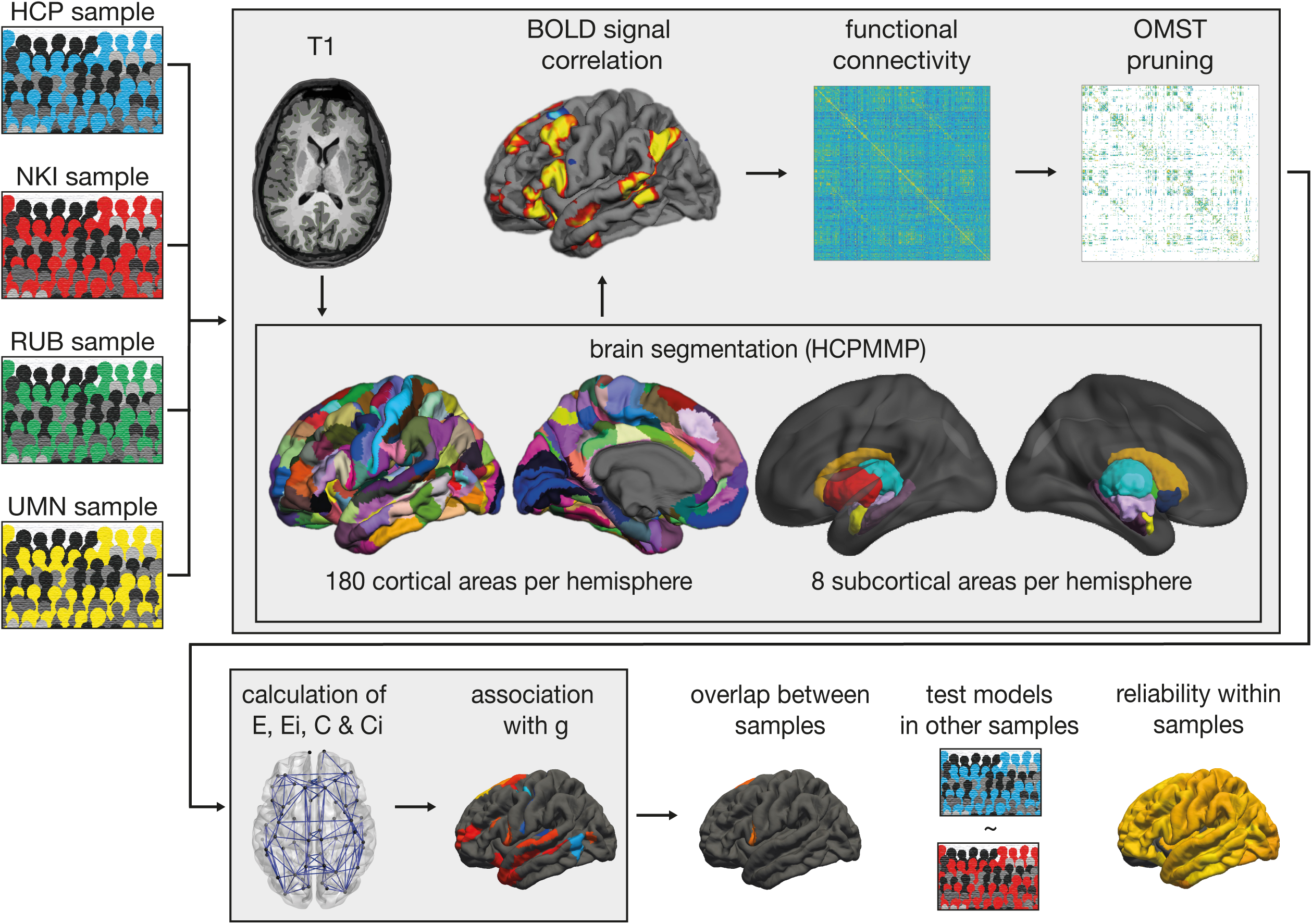
Pre-processing and analysis strategy of the four data sets. The HCP, NKI, RUB, and UMN samples were pre-processed in the same manner (light gray boxes). First, T1-weighted anatomical images were delineated into 180 cortical and 8 subcortical areas per hemisphere. Second, these areas were used as landmarks to extract mean time courses from resting-state images. Third, functional connectivity matrices were built by computing edge weights in the form of BOLD signal correlations. Fourth, pruning was applied to every data set to remove spurious connections from the network. Fifth, networks of all participants were pruned using OMST. Sixth, all functional connectivity matrices and graph theoretical metrics were computed. Seventh, partial correlation coefficients were calculated, and elastic-net regression was applied to investigate the association of graph theoretical metrics and *g* in all data sets. Eighth, the overlap between the results of the four different samples were compared and prediction models resulting from the elastic-net analysis in each sample were tested in the other independent samples. Finally, reliability of graph metrics was defined using the HCP and NKI samples.

### 2.4. Distribution of intelligence scores

The four samples included in our study employed different tests to assess intelligence. Hence, it is not possible to compare intelligence levels among samples directly. However, we utilized norming data of some tests to provide estimates for the intelligence levels of all samples as described in Stammen et al. ^[41]^. For the RUB data set, the I-S-T 2000 R subtests generated a mean IQ of 115 (SD = 13.0), one standard deviation above average. For the HCP data set, Dubois et al. ^[47]^ observed, based on norming data from the NIH toolbox subtests, that the sample’s mean scores for all tests were significantly higher than the general population means. Application of the standard Wechsler formulae revealed mean IQ scores of 114.1 (SD = 15.0) in the UMN sample and of 101.9 (SD = 13.1) in the NKI sample. The fact that three out of four samples had higher than average mean scores may have impacted the associations with functional brain metrics examined in our study.

### 2.5. Handedness measurement

Handedness was assessed using the Edinburgh Handedness Inventory ^[48]^. Here, participants answer ten questions regarding their preferred hand for everyday tasks, e.g. writing. The lateralization quotient (LQ) is defined as LQ = [(R – L)/(R + L)] x 100, with R indicating the sum of right-hand responses and L indicating the sum of left-hand responses. Handedness was coded 1 for all participants whose LQ was greater than or equal to 60 (right-handed) and 0 for all other participants (mixed- or left-handed). We chose this cut-off value based on Dragovic ^[49]^. Handedness was used as a control variable since the handedness ratios of our samples differed greatly.

### 2.6. Acquisition of anatomical data

#### 2.6.1. RUB Sample

Magnetic resonance imaging was conducted on a 3T Philips Achieva scanner with a 32-channel head coil. The scanner was located at Bergmannsheil University Hospital in Bochum, Germany. T1-weighted data were obtained by means of a high-resolution anatomical imaging sequence with the following parameters: MP-RAGE; TR = 8.179 ms; TE = 3.7 ms; flip angle = 8°; 220 slices; matrix size = 240 x 240; resolution = 1 mm x 1 mm x 1 mm; acquisition time = 6 minutes.

#### 2.6.2. HCP Sample

Magnetic resonance imaging was conducted on a Siemens 3T Connectome Skyra scanner with a 32-channel head coil and a “body” transmission coil designed by Siemens. The scanner was located at Washington University. T1-weighted data were obtained by means of a high-resolution anatomical imaging sequence with the following parameters: MP-RAGE; TR = 2400 ms; TE = 2.14 ms; flip angle = 8°; 256 slices; matrix size = 224 x 224; resolution = 0.7 mm x 0.7 mm x 0.7 mm; acquisition time = 7 minutes and 40 seconds.

#### 2.6.3. UMN Sample

All images were collected on a 3T Siemens Trio scanner at the Center for Magnetic Resonance Research at the University of Minnesota in Minneapolis, using a 12-channel head coil. T1-weighted data were obtained by means of a high-resolution anatomical imaging sequence with the following parameters: MP-RAGE; TR = 1900 ms; TE = 0.29 ms; flip angle = 9°; 240 slices; matrix size = 256 x 256; resolution = 1 mm x 1 mm x 1 mm; acquisition time = 8 minutes.

#### 2.6.4. NKI Sample

All images were collected on a 3T Siemens Trio scanner at the NKI in Orangeburg, New York, using a 32-channel head coil. T1-weighted data were obtained by means of a high-resolution anatomical imaging sequence with the following parameters: MP-RAGE; TR = 1900 ms; TE = 2.52 ms; flip angle = 9°; 176 slices; matrix size = 256 x 256; resolution = 1 mm x 1 mm x 1 mm; acquisition time = 4 minutes and 18 seconds minutes.

### 2.7. Acquisition of resting-state fMRI data

#### 2.7.1. RUB Sample

Functional MRI resting-state images were acquired using echo planar imaging (TR = 2000 ms, TE = 30 ms, flip angle = 90°, 37 slices, matrix size = 80 x 80, resolution = 3 mm x 3 mm x 3 mm, acquisition time = 7 minutes). Participants were instructed to lay still with their eyes closed and to think of nothing in particular.

#### 2.7.2. HCP *Sample*

For the HCP sample, four rsfMRI measurements were available (HCP_day_1_ses_1, HCP_day_1_ses_2, HCP_day_2_ses_1, and HCP_day_2_ses_2). On each of two consecutive testing days, two rsfMRI scans were acquired with opposite encoding directions, namely left-right (session 1) and right-left (session 2). Participants had their eyes open and fixated on a cross-hair. Images were acquired using echo planar imaging (TR = 720 ms, TE = 33 ms, flip angle = 52°, 72 slices, matrix size = 104 x 90, resolution = 2 mm x 2 mm x 2 mm, acquisition time = 14 minutes and 33 seconds).

#### 2.7.3. UMN *Sample*

Functional MRI resting-state images were acquired using echo planar imaging (TR = 2000 ms, TE = 28 ms, flip angle = 80°, 35 slices, matrix size = 64 x 64, resolution = 3.5 mm x 3.5 mm x 3.5 mm, acquisition time = 5 minutes). Participants performed a basic fixation task. They fixated on a cross-hair and pressed a button when the cross-hair changed colors (this happened five times). This was done to minimize eye movement and to ensure that participants stayed awake.

#### 2.7.4. NKI *Sample*

The NKI sample provided three rsfMRI measurements, namely NKI_TR645, NKI_TR1400, and NKI_TR2500. All of them were acquired on the same day. Participants had their eyes open and fixated on a cross-hair. Images were obtained using echo planar imaging with the following parameters. NKI_TR645: TR = 645 ms, TE = 30 ms, flip angle = 60°, 40 slices, matrix size = 74 x 74, resolution = 3 mm x 3 mm x 3 mm, acquisition time = 9 minutes and 46 seconds. NKI_TR1400: TR = 1400 ms, TE = 30 ms, flip angle = 65°, 64 slices, matrix size = 112 x 112, resolution = 2 mm x 2 mm x 2 mm, acquisition time = 9 minutes and 35 seconds. NKI_TR2500: TR = 2500 ms, TE = 30 ms, flip angle = 80°, 38 slices, matrix size = 72 x 72, resolution = 3 mm x 3 mm x 3 mm, acquisition time = 5 minutes and 5 seconds.

### 2.8. Imaging Processing

All data sets were processed in the same manner (see Figure 1).

#### 2.8.1. Anatomical Processing

Cortical surfaces of T1-weighted images were reconstructed using FreeSurfer (http://surfer.nmr.mgh.harvard.edu, version 6.0.0), along with the CBRAIN platform ^[50]^, following established protocols ^[51, 52]^. During pre-processing, we employed skull stripping, gray and white matter segmentation, and reconstruction and inflation of the cortical surface. These steps were done individually for each participant. We conducted slice-by-slice quality control and manually edited inaccuracies of automatic pre-processing. For brain segmentation, we used the Human Connectome Project’s multi-modal parcellation (HCPMMP), which comprises 180 areas per hemisphere and is based on structural, functional, topographical, and connectivity data from healthy participants ^[53]^. The original data provided by the Human Connectome Project were converted to annotation files matching the standard cortical surface in FreeSurfer (fsaverage). The fsaverage segmentation was transformed to each participant’s individual cortical surface and converted to volumetric masks. Additionally, we added eight subcortical gray matter structures per hemisphere to the parcellation (thalamus, caudate nucleus, putamen, pallidum, hippocampus, amygdala, accumbens area, ventral diencephalon) ^[54]^. Finally, six regions representing the four ventricles of the brain were defined to serve as a control for later BOLD signal analyses. All masks were linearly transformed into the native space of the resting-state images and used as landmarks for graph-theoretical connectivity analyses.

#### 2.8.2. Resting-state fMRI Processing

Pre-processing of rsfMRI data was conducted with the FSL toolbox MELODIC. The first two images of each resting-state scan were discarded to ensure signal equilibration, motion and slice timing correction, and high-pass temporal frequency filtering (0.005 Hz). We did not apply spatial smoothing so the introduction of spurious correlations in neighboring voxels would be avoided. All 376 brain regions (360 cortical and 16 subcortical) were converted into the native space of the resting-state images for functional connectivity analysis. For each region, we computed a mean resting-state time course by averaging the time courses of relevant voxels. Partial correlations between the average time courses of all cortical and subcortical regions were calculated, while controlling for all six motion parameters as well as average time courses extracted from white matter regions and ventricles ^[55]^.

### 2.9. Statistical Analysis

#### 2.9.1. Computation of graph theoretical metrics

All analyses were carried out using MATLAB version R2021b (The MathWorks Inc., Natick, MA). For graph-theoretical connectivity analyses, we defined a brain network with 376 nodes, consisting of the 360 cortical and 16 subcortical regions described above. Since each brain region can theoretically be connected to each other brain region, there was a total of 70500 edges (one matrix triangle of a symmetrical 376-by-376 adjacency matrix without self-connections on the diagonal). Each edge was weighted based on the BOLD signal correlation between the two brain regions at the edge’s endpoints. Negative correlation coefficients were replaced with zeros. The four rsfMRI scans of the HCP sample were treated as individual data sets (HCP_day_1_ses_1, HCP_day_1_ses_2, HCP_day_2_ses_1, and HCP_day_2_ses_2) but also conflated into bigger data sets. In more detail, we computed functional connectivity matrices for HCP_day_1, HCP_day_2, and HCP_day_1_day_2. Relevant BOLD signal correlations were calculated after concatenating time courses for the two scans on each day for HCP_day_1 and HCP_day_2 and after concatenating time courses for all four scans for HCP_day_1_day_2. For the sake of readability, we chose to only report results from HCP_day_1_ses_1 and HCP_day_1_ses_2, while omitting the remaining individual data sets, which produced highly similar results. To ensure that correlation coefficients were normally distributed, we Fisher z-transformed all values ^[56]^. To prevent our analyses being affected by spurious connections ^[55, 57]^, while also avoiding arbitrary graph thresholds, we applied a data-driven topological filtering to the individual functional connectivity matrices using orthogonal minimum spanning trees (OMST) ^[58, 59]^. Graph metrics were computed using the Brain Connectivity Toolbox ^[30]^. We chose global efficiency, nodal efficiency, global clustering, local clustering, and small-world propensity as measures to quantify network connectivity.

##### 2.9.1.1. Orthogonal minimum spanning trees

For network pruning we used the Topological Filtering Toolbox as provided and described by Dimitriadis et al. ^[58, 59]^ (https://github.com/stdimitr/topological_filtering_networks). The OMST approach focuses on optimizing a function of the global efficiency (see below) and the wiring cost of the network (ratio of the total weight of the selected edges, aggregated over multiple iterations of OMST, of the filtered graph to that of the original fully weighted graph), while ensuring that the network is fully connected. It is a data-driven, iterative procedure creating multiple minimum spanning trees (MST) that are orthogonal to each other. MST creates a subgraph with the minimum set of edges (N-1) that connect all N nodes using Kruskal’s algorithm. After extracting the first MST, its N-1 edges are zeroed out in the original graph to maintain orthogonality and a second MST connecting all the N nodes with minimal total distance is extracted. These steps are repeated, and connections are aggregated across OMST until optimization of the function Global Cost Efficiency = global efficiency – wiring cost. The final filtered graph includes all MST that have been removed from the original graph. OMST is applied to every participant and the trade-off is optimized for each network individually, leaving all participants with their unique, but fully connected network. Dimitriadis et al. ^[59]^ have shown that the OMST algorithm generated sparse graphs that outperformed many other thresholding schemes based on recognition accuracy and reliability of network metrics. OMST has been successfully applied in resting-state functional connectivity research^[60]^.

##### 2.9.1.2. Global and nodal efficiency

Efficiency is a measure of functional integration capturing the brain’s ability to exchange specialized information within a network of distributed brain regions ^[30]^. Shorter path lengths, indicating that two nodes are connected by a path with few edges, generally allow more efficient communication since signal transmission in fewer steps affords less opportunity for noise, interference or attenuation ^[3]^. The shortest paths lengths 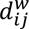 between all pairs of nodes can be obtained by calculating the inverse of the weighted adjacency matrix and running a search algorithm such as Dijkstra’s ^[61]^ implemented in the Brain Connectivity Toolbox ^[30]^. The global efficiency (*E*) of a network is calculated as the average inverse shortest path length between each pair of nodes *i* and *j* within the network *G* ^[1]^. On the level of single nodes, nodal efficiency (*E_i_*) can be determined as a metric quantifying the importance of each specific node *i* for information transfer within the network ^[32]^. It is calculated as the average inverse shortest path length between a particular node and all other nodes within the network, where n is the number of nodes ^[1, 30]^.

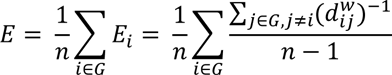

##### 2.9.1.3. Global and local clustering

Clustering is a measure quantifying the extent of local “cliquishness” and serves as an indicator of local connection segregation ^[3, 7, 32]^. The local clustering coefficient *C_i_* represents the probability that two randomly selected neighbors of a particular node *i* are also neighbors of each other (i.e. connections among these three nodes form a triangle). *C_i_* is computed by dividing the existing connections among the node’s neighbors by all possible connections ^[7, 32]^. The global clustering coefficient *C* is calculated as the average of all clustering coefficients of all nodes and reflects the occurrence of clustered connectivity around single nodes ^[30]^. In the context of neurobiology, clustered connectivity indicates the existence of brain regions that constitute functionally coherent units with a high degree of within-unit information exchange ^[3]^.

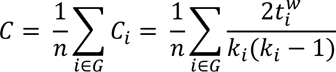

Here, n is the number of nodes, *k_i_* is the number of edges connected to a node *i*, and *t_i_* is the number of triangles attached to a node *i* ^[7, 30, 62]^. As required by the Brain Connectivity Toolbox, coefficients were normalized to be between 0 and 1 before calculation ^[30]^.

##### 2.9.1.4. Small-world propensity

Small-world networks are characterized by high local clustering (segregated functionally coherent units) and short path lengths among clusters. The latter is typically realized by robust numbers of integrating, intermodular, long-range links ^[6, 7, 30]^. To quantify the small-world structure of participants’ functional connectivity matrices, we used the small-world propensity function as proposed by Muldoon et al. ^[63]^. This function reflects the discrepancy of a network’s clustering coefficient, *C_obs_*, and characteristic path length, *L_obs_*, from comparable lattice (*C_latl_*, *L_latt_*) and random (*C_rand_*, *L_rand_*) networks ^[63]^.

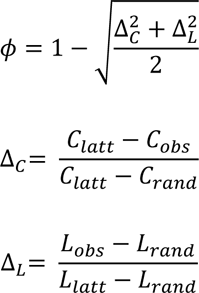

#### 2.9.2. Partial correlations

The following statistical analyses were conducted in R Studio (1.3.1093) with R version 4.1.0. (2021-05-18). For each sample, outlier control was conducted, and outliers were winsorized ^[46]^. Data points were treated as outliers if they deviated more than three interquartile ranges from the relevant variable’s group mean (*g*, global efficiency, global clustering, small-world propensity) and replaced by the respective threshold.

We calculated partial correlations between *g* and the following network properties: global efficiency, global clustering, and small-world propensity. We used the partial.cor function as implemented in the RcmdrMisc package to obtain two-sided *p*-values corrected for multiple comparisons using Holm’s method ^[64]^. Control variables were age, sex, age*sex, age^2^, age^2^*sex, and handedness.

#### 2.9.3. Brain area specific analysis via elastic-net regression

To examine the associations between general intelligence and network properties of single brain areas, we employed regularized elastic-net regression. This approach does not use *p*-values to determine the statistical significance of a predictor. Hence, it does not require a standard correction procedure for multiple comparisons (e.g. False Discovery Rate or Bonferroni). Instead, it utilizes regularization and cross-validation permitting models with many independent variables in a small sample. Regularization puts a penalty on all effect sizes. The penalty is calculated using k-fold cross-validation, a mechanism to avoid overfitting. This leads to small effect sizes being regularized to zero. Every non-zero effect size contributes uniquely to the association. In previous studies, elastic-net regression has already been successfully applied to research questions concerned with the association between brain properties and intelligence ^[65–67]^.

For this step, we used the cv.glmnet function from the glmnet package. Alpha was set to 0.5, k was set to 10. All variables were standardized and residualized for age, sex, age*sex, age^2^, age^2^*sex, and handedness. We computed two elastic-net regression models for every rsfMRI measurement from all samples (RUB, HCP_day_1, HCP_day_1_ses_1, HCP_day_1_ses_2, HCP_day_2, HCP_day_1_day_2, UMN, NKI_TR645, NKI_TR1400, NKI_TR2500). The dependent variable was always *g*, the independent variables were either nodal efficiency of all 376 network nodes or local clustering of all 376 network nodes.

#### 2.9.4. Overlap among samples and result validity

To assess result validity for the elastic-net regression analysis, we identified brain areas whose properties emerged as relevant associates of *g* among multiple regression models from all data sets. To this end, we included the elastic-net results from the RUB, HCP_day_1_day_2, UMN, and NKI_TR645 samples. HCP_day_1_day_2 was chosen to ensure comparability with previous studies ^[47, 68]^. NKI_TR645 was chosen over NKI_TR1400 and NKI_TR2500 since its TR is closer to that of the HCP sample. Furthermore, we investigated the overlap with results obtained from HCP data, once for the two measurement days (HCP_day_1 and HCP_day_2) and once for the two measurement sessions of the first day (HCP_day_1_ses_1 and HCP_day_1_ses_2). We also analyzed the intrasession overlap of results obtained from NKI data. Finally, we compared the two TR = 2000 ms data sets RUB and UMN as well as the two TR ≤ 720 ms data sets HCP and NKI.

As elastic-net regression is a data-driven procedure, relatively small fluctuations can influence which effects will remain non-zero after regularization. Thus, the comparison of elastic-net results alone is not sufficient to judge the validity of the models. To judge validity, we tested the models defined by elastic-net in each sample in all other samples. In more detail, we calculated multiple linear regression analyses in the testing samples with the predictors identified in one data set serving as independent variables and *g* serving as dependent variable. For example, predictors identified in the HCP_day_1_day_2 sample would be used to predict *g* in RUB, UMN and NKI_TR645. Control variables were age, sex, age*sex, age^2^, age^2^*sex, and handedness. We used the coefficient of determination along with its significance value to conclude whether the set of predictors identified in one sample predicted intelligence successfully in the other samples. We used the elastic-net results of each sample in all other samples once. However, due to space limitations we will only show the results of four samples (RUB, HCP_day_1_day_2, UMN, and NKI_TR645) in the results section. For these analyses, the *p*-value was corrected for multiple comparisons using Bonferroni ^[69]^.

#### 2.9.5. Test-retest reliability of graph metrics

The HCP and NKI samples afforded examination of the graph theoretical measures’ test-retest reliability. In the HCP data set, we assessed reliability once between measurement days (HCP_day_1 and HCP_day_2) as well as within the first measurement day (HCP_day_1_ses_1 and HCP_day_1_ses_2). In the NKI data set, we examined the reliability of measurement sessions (NKI_TR645, NKI_TR1400, and NKI_TR2500). First, we investigated the reliability of global metrics (global efficiency, global clustering, and small-world propensity). We computed an intra-class correlation (ICC) (3,1) two-way mixed effect model ^[70]^, which is a robust measure of reliability in fMRI research ^[71]^. Second, we investigated the reliability of nodal metrics (nodal efficiency and local clustering) for every brain area independently.

## 3. Results

### 3.1. Association between global brain metrics and *g* (pruned across all data sets)

Table 1 shows partial correlations between general intelligence and three global brain metrics (global efficiency, global clustering, and small-world propensity) controlling age, sex, age*sex, age^2^, age^2^*sex, and handedness. Overall, we did not find consistent significant associations between any global brain metric and general intelligence. Nevertheless, there were significant positive associations between general intelligence and global clustering in the HCP sample (day 1 session 1 and day 1 session 2) as well as in NKI_TR645 and between general intelligence and small-world propensity in the HCP sample (day 1 session 1 and day 1 session 2) as well as in NKI_TR645 and in NKI_TR1400. All significant effect sizes were small with mean *r* = .13 (SD = .03) ^[72]^.

**Table 1.** Partial correlation results between general intelligence and global brain metrics.

| Data set | Global efficiency |  | Global clustering |  | Small-world propensity |  |
| --- | --- | --- | --- | --- | --- | --- |
|  | *r* | *p* | *r* | *p* | *r* | *p* |
| HCP_day_1 | -.03 | .298 | .04 | .173 | .01 | .666 |
| HCP_day_1_ses_1 | .01 | .814 | .12 | **<.001** | .09 | **.004** |
| HCP_day_1_ses_2 | -.04 | .201 | .10 | **.001** | .14 | **<.001** |
| HCP_day_2 | -.04 | .212 | .04 | .221 | .01 | .693 |
| HCP_day_1_day_2 | -.01 | .779 | -.04 | .223 | -.00 | .958 |
| NKI_TR645 | -.10 | .052 | .13 | **.011** | .19 | **<.001** |
| NKI_TR1400 | -.10 | .067 | .09 | .091 | .16 | **.003** |
| NKI_TR2500 | -.09 | .102 | .03 | .601 | -.08 | .143 |
| RUB | .07 | .118 | .05 | .230 | .02 | .731 |
| UMN | -.03 | .656 | .00 | .606 | .05 | .390 |

*Note.* Partial correlations between general intelligence and global efficiency, global clustering, and small-world propensity. Control variables were age, sex, age\*sex, age<sup>2</sup>, age<sup>2</sup>\*sex, and handedness. *p*-values < .05 are in boldface.

### 3.2. Association validity for nodal efficiency and *g*

Figure 2 depicts the associations between general intelligence and nodal efficiency in all data sets (left and middle columns) as well as the overlaps between data-set pairs (right column). A complete list of all cortical and subcortical areas exhibiting unique associations with general intelligence is in Supplementary Table S1. Figure 2 does not show overlap among all data sets (RUB, HCP_day_1_day_2, UMN, and NKI_TR645) since there were no brain areas that had non-zero effect sizes in all data sets. Table 2 shows the percentage of areas identified in elastic-net analysis that overlapped between data sets depicted in Figure 2. Even when comparing different measurement sessions from the same data set (HCP_day_1 and HCP_day_2, HCP_day_1_ses_1 and HCP_day_1_ses_2, and NKI_TR645, NKI_TR1400 and, NKI_TR2500), no more than 18.5% of brain regions overlapped (see Supplementary Table S2 for a complete list of overlapping areas).

**Figure 2.**
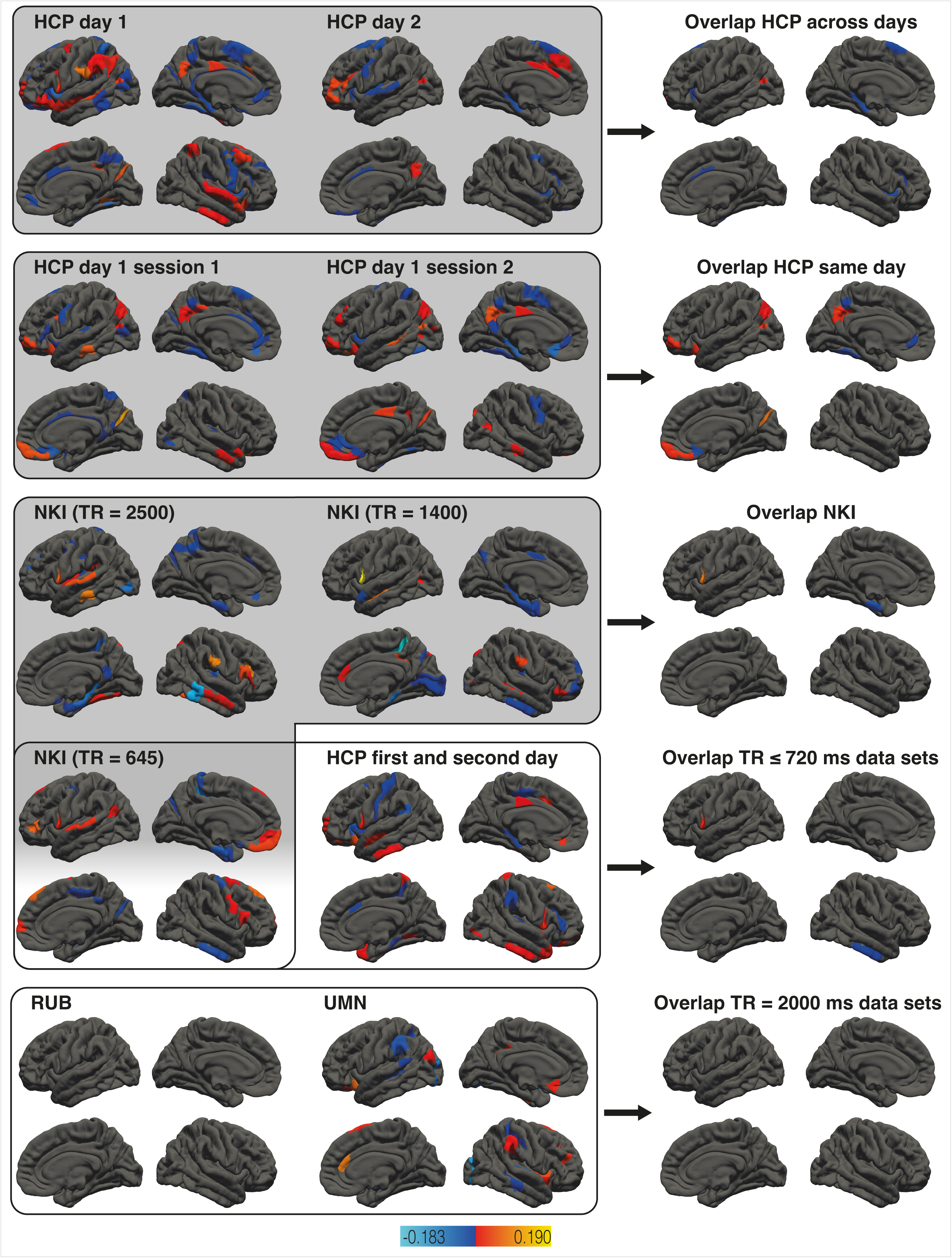
Association between nodal efficiency and general intelligence. The left and middle columns show the results of brain area specific analyses relating nodal efficiency and general intelligence in all data sets. Brain areas exhibiting relevant associations are color-coded based on effect size. The overlaps between framed data-set pairs (left and middle columns) are depicted next to them (right column). Effect sizes of overlapping areas are averaged. Rows with gray backgrounds show within-sample overlaps, rows with white backgrounds between-sample overlaps. Subcortical areas and hemispheric specific illustrations are not shown but listed in Supplementary Table S2.

**Table 2.** Number of areas associated with intelligence in each data set and overlap of areas associated with intelligence (nodal efficiency) between data sets.

| Data set 1 | Data set 2 | Data set 3 | Total | Overlapping areas (%) |
| --- | --- | --- | --- | --- |
| HCP_day_1: 91 | HCP_day_2: 33 | - | 108 | 16 (14.8%) |
| HCP_day_1_ses_1: 56 | HCP_day_1_ses_2: 53 | - | 92 | 17 (18.5%) |
| NKI_TR645: 28 | NKI_TR1400: 31 | NKI_TR2500: 44 | 101 | 2 (2%) |
| RUB: 0 | UMN: 31 | - | 31 | 0 (0%) |
| NKI_TR645: 28 | HCP_day_1_day_2: 40 | - | 66 | 2 (3%) |

*Note.* Total = sum of areas associated with intelligence of data set 1, 2 (and 3) minus the number of overlapping areas. The number of overlapping areas relates to the mere number of areas, irrespective of the direction of association.

Table 3 shows how well the models defined via elastic-net regression in four samples (HCP_day_1_day_2, NKI_TR645, RUB, and UMN) tested in the other samples via multiple regression analyses performed. After correction for multiple comparisons using Bonferroni ^[69]^, the predictors identified in one sample did not significantly predict general intelligence in other samples. The explained variance in general intelligence varied between 3% and 16% in the different samples, while it was up to 40% when the predictors were tested with multiple regression analysis in the same sample. The complete results including all identified predictor sets tested in all samples via multiple linear regression analyses (uncorrected for multiple comparisons) for nodal efficiency can be found in Supplementary Tables S6 and S7. Supplementary Table S6 shows the coefficients of determination and the uncorrected *p*-values, while Supplementary Table S7 depicts the number of significant predictors when tested in other samples. The extent of explained variance in general intelligence predicted using a set of nodal efficiency variables identified in a different sample varied between 3% and 41% (see Table S6). 2% to 30% of the predictors identified via elastic-net were significantly associated with general intelligence in the new sample (uncorrected for multiple comparisons, see Supplementary Table S7).

**Table 3.** Results of multiple linear regression analyses for nodal efficiency. Areas identified via elastic-net in one sample (left column) have been used as predictors for general intelligence in the other samples.

|  | HCP_day_1_day_2 |  | NKI_TR645 |  | RUB |  | UMN |  |
| --- | --- | --- | --- | --- | --- | --- | --- | --- |
|  | R<sup>2</sup> | *p* | R<sup>2</sup> | *p* | R<sup>2</sup> | *p* | R<sup>2</sup> | *p* |
| HCP_day_1_day_2<br>(40 predictors) | .17 | **<.001** | .15 | .022 | .10 | .034 | .16 | .302 |
| NKI_TR645<br>(28 predictors) | .03 | .277 | .28 | **<.001** | .07 | .083 | .12 | .253 |
| RUB<br>(0 predictors) | - | - | - | - | - | - | - | - |
| UMN<br>(31 predictors) | .04 | .136 | .11 | .105 | .05 | .707 | .40 | **<.001** |

*Note.* Multiple linear regression analyses predicting general intelligence in other samples based on areas with non-zero effect sizes identified via elastic-net in one sample. Control variables were age, sex, age\*sex, age<sup>2</sup>, age<sup>2</sup>\*sex, and handedness. R<sup>2</sup> = coefficient of determination. Multiple comparisons have been corrected using Bonferroni. *p*-values < .003 (= .05/16) are in boldface.

### 3.3. Association validity among local clustering and *g*

Figure 3 depicts the associations between general intelligence and local clustering in all data sets (left and middle columns) as well as result overlaps between data-set pairs (right column). See Supplementary Table S3 for a complete list of all cortical and subcortical areas exhibiting unique associations with general intelligence. There was no area that had a non-zero effect size in all four data sets. As with nodal efficiency, local clustering did not yield strongly overlapping results even when comparing results from the same data set but different measurement sessions (HCP_day_1 and HCP_day_2, HCP_day_1_ses_1 and HCP_day_1_ses_2, and NKI_TR645, NKI_TR1400, and NKI_2500). Here, no more than 10% of brain areas overlapped (see Fig. 3 and Table 4). For a complete list of overlapping areas see Supplementary Table S4.

**Figure 3.**
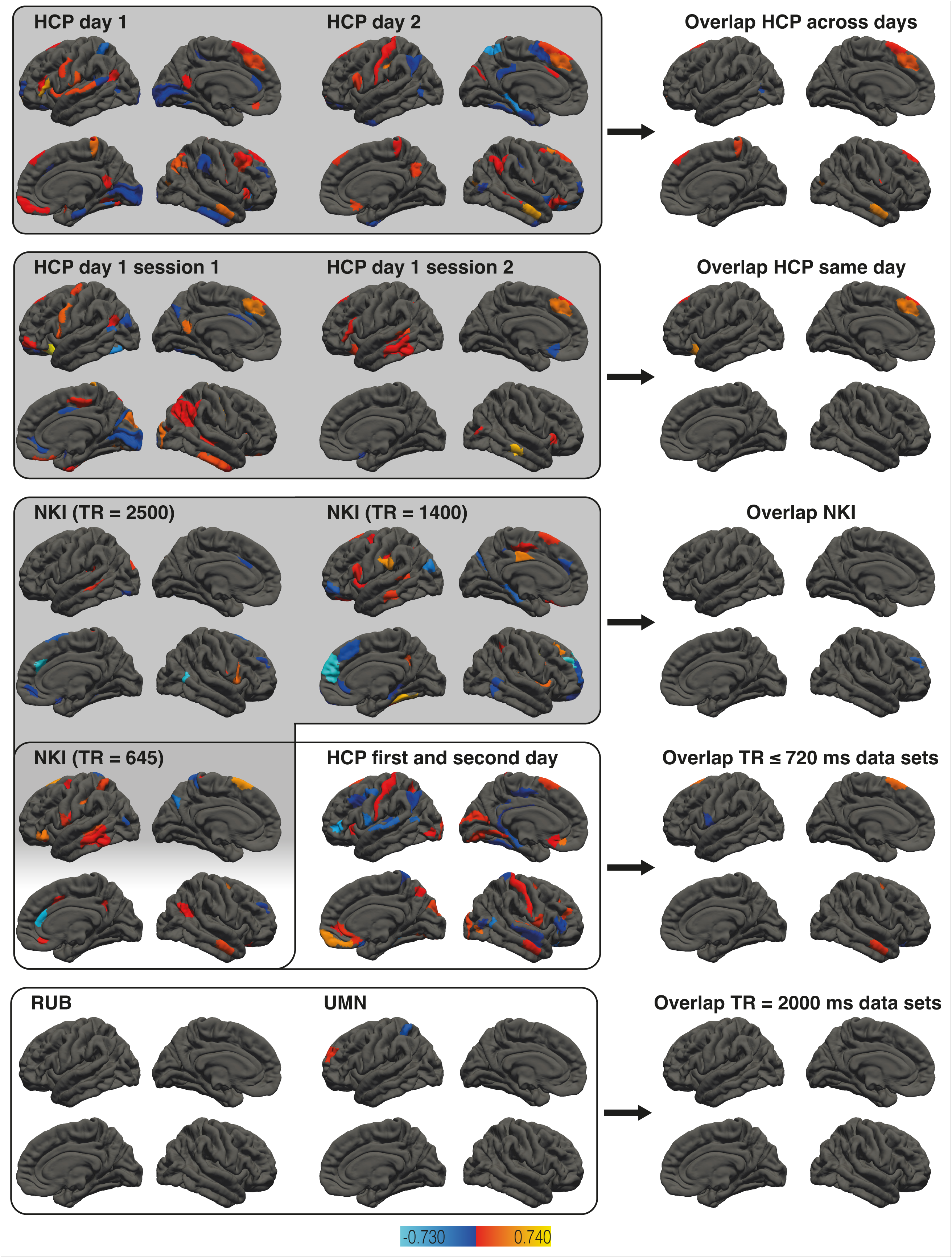
Association between local clustering and general intelligence. The left and middle columns show the results of brain area specific analyses relating local clustering and general intelligence in all data sets. Brain areas exhibiting relevant associations are color-coded based on effect size. The overlap between framed data set pairs (left and middle columns) are depicted next to them (right column). Effect sizes of overlapping areas are averaged. Rows with gray backgrounds show within-sample overlaps, rows with white backgrounds between-sample overlaps. Subcortical areas and hemispheric specific illustrations are not shown but listed in Supplementary Table S3.

**Table 4.**
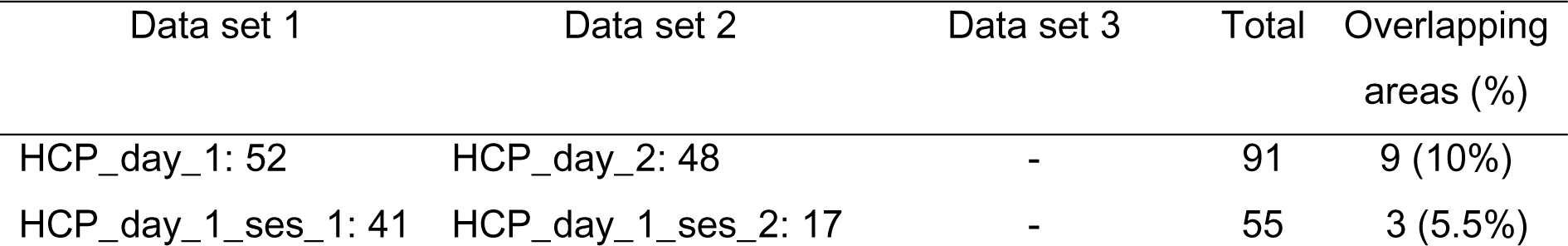
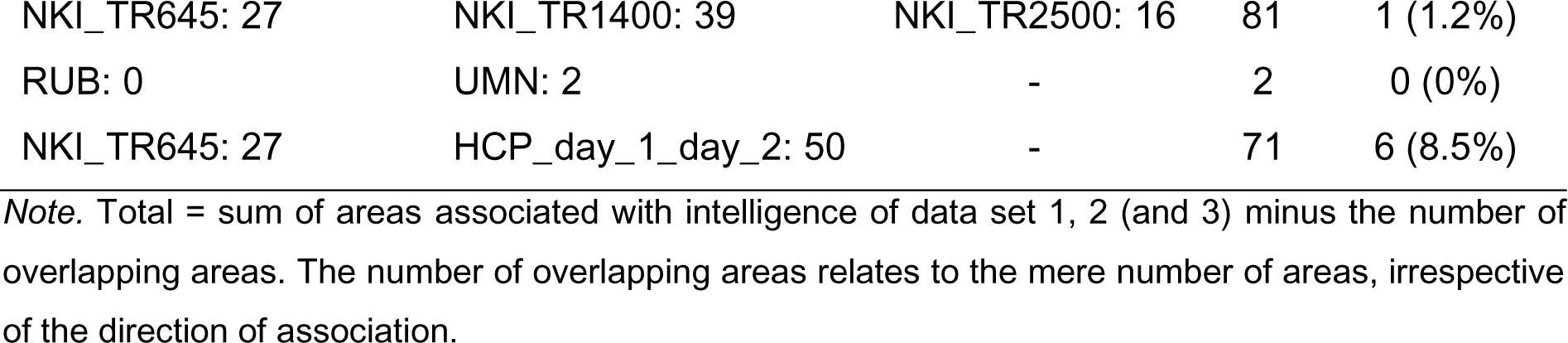
Number of areas associated with intelligence in each data set and overlap of areas associated with intelligence (local clustering) between data sets.

Table 5 shows the performance of the models defined via elastic-net regression in four samples (HCP_day_1_day_2, NKI_TR645, RUB, and UMN) in the other samples tested with multiple regression analyses. After correction for multiple comparisons using Bonferroni ^[69]^, the predictor set of the HCP_day_1_day_2 sample significantly predicted general intelligence in the NKI_TR645 sample and vice versa, explaining 5% to 21% of variance in *g*. In general, the explained variance in general intelligence varied between 0% and 21% in the different samples. Only using the predictors identified in the elastic-net regression in the NKI_TR645 sample for multiple linear regression analysis in the same sample resulted in a higher explained variance (27%). The complete results including all identified predictor sets tested in all samples via multiple linear regression analyses (uncorrected for multiple comparisons) for local clustering can be found in Supplementary Tables S8 and S9. Supplementary Table S8 shows the coefficients of determination and the uncorrected *p*-values, while Supplementary Table S9 depicts the number of significant predictors when tested in other samples. The extent of explained variance in general intelligence predicted using a set of local clustering variables identified in a different sample varied between 0 and 22% (see Supplementary Table S8). 0% to 50% (25%, when ignoring the UMN sample with only two predictors) of the predictors identified via elastic-net were significantly associated with general intelligence in the new sample (uncorrected for multiple comparisons, see Supplementary Table S9).

**Table 5.**
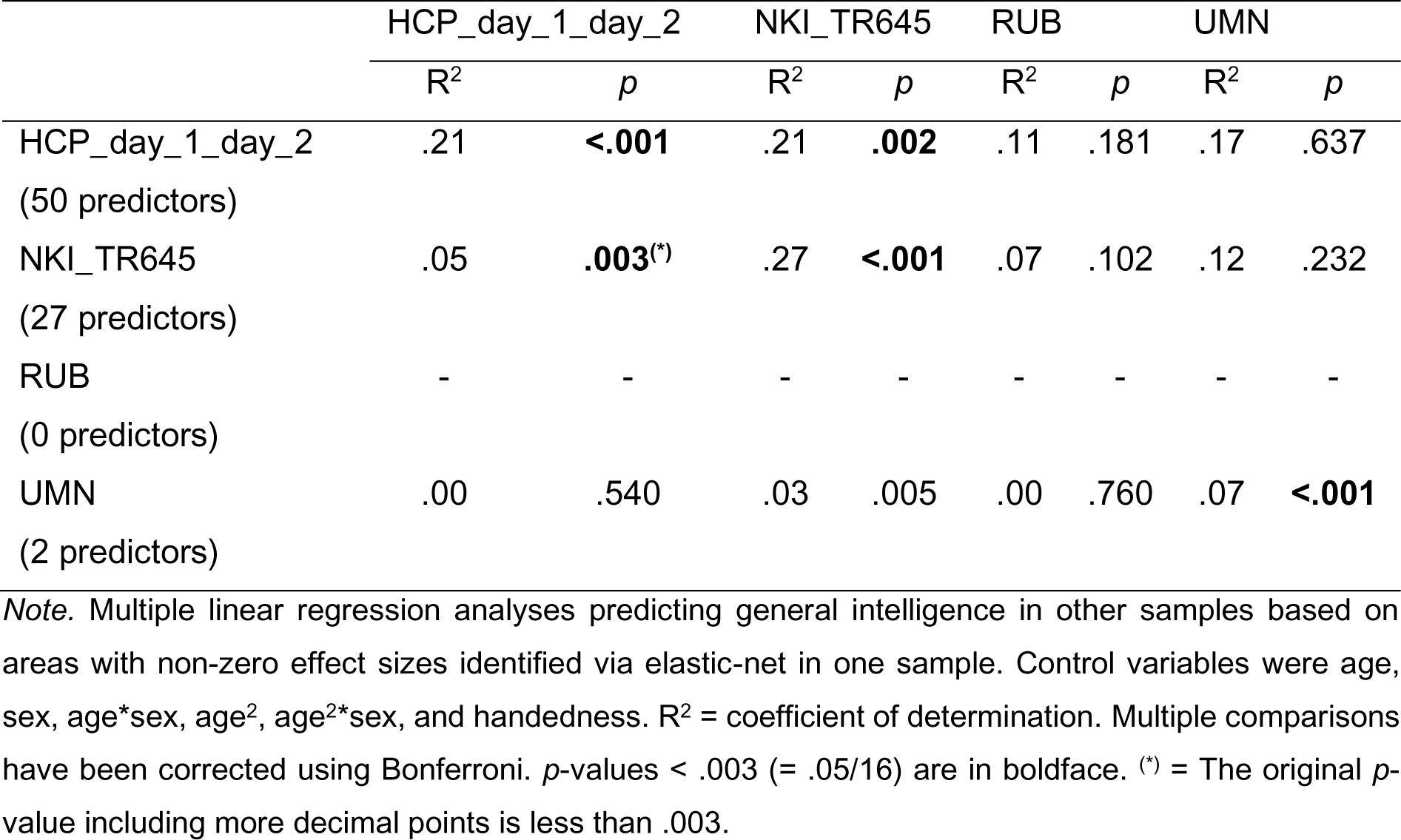
Results of multiple linear regression analyses for local clustering. Areas identified via elastic-net in one sample (left column) have been used as predictors for general intelligence in the other samples.

### 3.4. Global brain metric reliability

Global graph metric reliability was quantified using ICC. Relevant coefficients are listed in Table 6. Global efficiency reliability across days can be considered moderate to fair ^[70, 73]^. Same-day reliability can be considered good in the NKI sample and good to excellent in the HCP sample. In contrast, global clustering reliability can be considered only poor to fair across days. Same-day reliability can be considered moderate to fair for the HCP sample and poor to fair in the NKI sample. The same applies to small-world propensity reliability.

**Table 6.** ICC of global brain metrics.

| Data set | Global efficiency |  | Global clustering |  | Small-world propensity |  |
| --- | --- | --- | --- | --- | --- | --- |
|  | *r* | *p* | *r* | *p* | *r* | *p* |
| HCP across days | .54 | < .001 | .43 | < .001 | .47 | < .001 |
| HCP same day | .75 | < .001 | .53 | < .001 | .54 | < .001 |
| NKI | .63 | < .001 | .47 | < .001 | .46 | < .001 |

### 3.5. Reliability of nodal efficiency and local clustering

Figure 4 depicts reliability of nodal efficiency and local clustering as quantified by ICC. Reliability varied greatly among areas, with e.g. orbitofrontal and anterior cingulate areas showing relatively poor reliability. ICC for nodal efficiency in the HCP sample across days was between .61 and .30, with 37% of values being higher than .50 (moderate to fair reliability). ICC for nodal efficiency in the HCP sample on the same day was between .75 and .29 (ICC in 88% of areas > .50). ICC for nodal efficiency in the NKI sample was between .63 and .14 (ICC in 43% of areas > .50). ICC for local clustering in the HCP sample across days was between .51 and .10 (ICC in 0.2% of areas > .50). ICC for local clustering in the HCP sample on the same day was between .51 and .03 (ICC in 0.2% of areas > .50). ICC for local clustering in the NKI sample was between .43 and .01.

**Figure 4.**
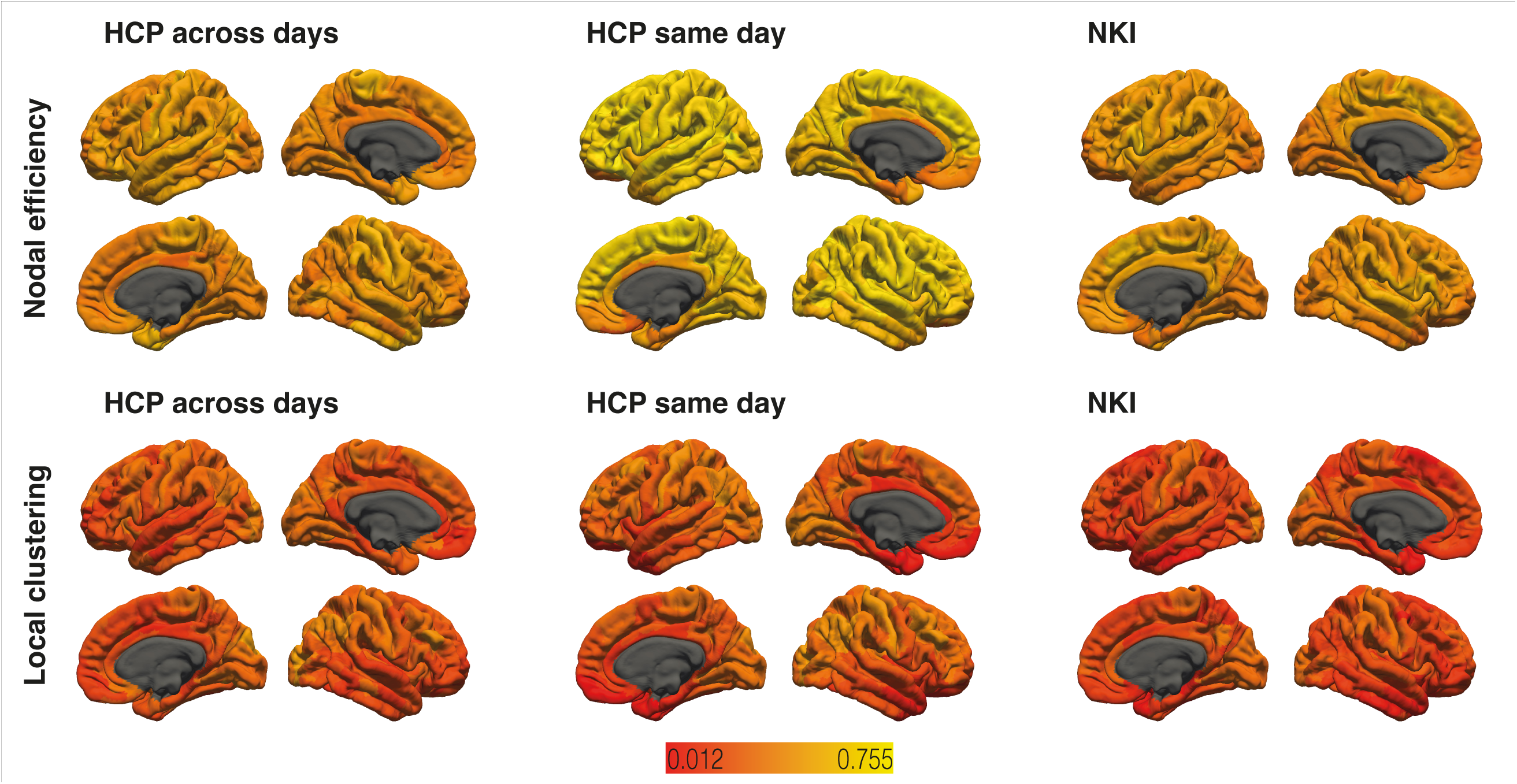
Reliability of nodal efficiency and local clustering. Intra-class correlation coefficients (ICC) of areas where the calculation was possible are color-coded with red indicating low ICC and yellow indicating high ICC.

## Discussion

While van den Heuvel et al. ^[21]^ found evidence for association between general intelligence and global brain functional network efficiency, newer observations question the proposed relation ^[36–38]^. The primary goal of our study was to investigate whether replicable associations between general intelligence and functional network properties exist using a multi-center approach. We computed partial correlations to examine the associations between *g* and global brain efficiency, global clustering, and small-world propensity and we employed regularized elastic-net regression analysis to examine the associations between general intelligence and the two region-specific metrics nodal efficiency and local clustering. On the global level, general intelligence showed no significant associations with global efficiency in any data set, but significant positive associations with global clustering coefficient as well as with small-world propensity in two data sets. On the level of single brain regions, there were no brain areas that consistently showed positive or negative non-zero effect sizes across all data sets for nodal efficiency or local clustering. We found the overlap of elastic-net results to be poor when comparing different data sets. Moreover, comparing different measurements from the same participants (HCP across days, HCP same day, and NKI) did not result in an overlap stronger than 18.5%. Using the areas identified via elastic-net regression in one sample to predict general intelligence via multiple linear regression in other samples was not successful regarding nodal efficiency. Regarding local clustering, areas associated with general intelligence in the HCP sample were able to predict general intelligence in the NKI sample and vice versa. Reliability estimates of the global and region-specific graph metrics were mostly fair to moderate, but poor for local clustering.

Our observation that general intelligence exhibited no significant association with global efficiency in any data set is consistent with results reported by previous studies ^[36–38]^. It strengthens Kruschwitz et al.’s ^[38]^ position that there is no robust relation between general intelligence and the efficiency of the brain’s intrinsic functional architecture. Although they employed multiple network definition schemes, ranging from 21637 nodes to 100 nodes, they did not find any significant association between global efficiency and a composite score of cognitive performance in the HCP data set. There were several differences between our study and theirs. We used different approaches for data pre-processing and brain network definition, we employed pruning via OMST instead of thresholding the networks based on connection strength, and we estimated *g* instead of using a composite score based on subtests from the NIH Toolbox. Despite the differences between the study designs, we also found no association, neither in the HCP data set nor in the other data sets. Furthermore, reliability of global efficiency was overall moderate to good, thus lack of reliability is unlikely to be the only reason for the non-significant results. Overall, our study has provided further evidence that there is no robust association between global efficiency and general intelligence.

We observed significant positive associations between general intelligence and global clustering coefficient in two HCP recordings, namely HCP_day_1_ses_1, HCP_day_1_ses_2, and NKI_TR645. Intelligence seemed, in those data sets, to involve functional brain architecture characterized by clustered connectivity around single nodes. A possible explanation for this might be that connected brain regions exhibit high information exchange and constitute functionally coherent units ^[3]^, enabling faster and/or more direct information processing. However, this result could not be considered robust given that it could not be detected in the other data sets. Van den Heuvel et al. ^[21]^, Pamplona et al. ^[36]^, and Kruschwitz et al. ^[38]^ also all reported non-significant results. Surprisingly, while the global clustering coefficients of the recordings HCP_day_1_ses_1 and HCP_day_1_ses_2 was associated with general intelligence, this effect did not appear in the combined HCP_day_1 sample. This could be due to random fluctuations, but another possible reason could be the poor test-retest reliability of the global clustering coefficient (see Table 6). It is noteworthy that we constructed highly individualized functional connectomes via OMST which have been shown to have a greater test-retest reliability compared to conventional thresholds ^[74]^. Improvements in test-retest reliability could be achieved by using whole-brain parcellations based on models integrating local gradient and global similarity approaches (e.g. Schaefer atlas ^[75]^) or focusing on higher frequencies in the slow band for time series filtering to derive the connectivity ^[74]^. It is considered well established that the human brain exhibits a small-world organization ^[1, 6–8]^. We observed significant positive associations between general intelligence and small-world propensity in two data sets, namely HCP (HCP_day_1_ses_1 and HCP_day_1_ses_2 recordings) and NKI (NKI_TR645 and NKI_TR1400 recordings). The combination of a high level of local clustering and robust long-distance connections ^[1, 6–8]^ seemed to be beneficial for general intelligence. However, as for global clustering coefficient, our result could not be considered robust as not all data sets revealed a significant positive association. However, failing to show this association in the smaller samples might be due to other reasons such as lack of power (e.g. UMN) or lack of test-retest reliability (e.g. HCP_day_1).

Results from the elastic-net analysis of specific brain regions showed rather low overlap among different samples. When comparing the regularized elastic-net regression results from the RUB, HCP, UMN, and NKI data sets, we did not find a single brain area that exhibited consistent non-zero associations with nodal efficiency or local clustering. Surprisingly, overlaps across days and even on the same day from the same data set were also rather poor. We found the maximum proportion of overlapping areas (irrespective of direction of association) to be 18.5% for nodal efficiency (see Table 2) and 10% for local clustering (see Table 4).

Elastic-net regression is a data-driven procedure, meaning that relatively small fluctuations can influence which effects will remain non-zero after regularization. Therefore, we tested the models defined by elastic-net in the four samples (HCP_day_1_day_2, NKI_TR645, RUB, and UMN) in all other samples via multiple linear regression analysis. There was no successful prediction of general intelligence based on nodal efficiency (see Table 3). Although the predictor set of local clustering areas of the HCP_day_1_day_2 data set significantly predicted general intelligence in the NKI_TR645 data sets and vice versa, there was no successful prediction in the UMN or RUB data set (see Table 5). Thus, there was no predictor set that successfully predicted general intelligence in all other data sets, neither for nodal efficiency nor for local clustering. One possible reason for the lack of robustness could be that the graph theoretical measures used in our study are not reliable. Reliability of nodal efficiency was overall fair to moderate, making it unlikely that low reliability is the only reason for the lack of robustness. However, reliability of local clustering was mostly poor. Thus, even though the predictors associated with general intelligence in HCP_day_1_day_2 also predicted general intelligence in NKI_TR645 and vice versa, these results should be interpreted with caution. Overall, our results indicate that resting-state functional connectivity nodal efficiency and local clustering do not reliably predict general intelligence.

However, the predictor set identified in one sample via elastic-net regression explained on average (uncorrected for multiple comparisons) 13.3% of variance in general intelligence in another sample for nodal efficiency and 8.7% of variance in general intelligence in another sample for local clustering (see Supplementary Tables S6 and S8). Supplementary Tables S7 and S9 reveal that only a small proportion of the predictors identified in the elastic-net regression also remained significant (uncorrected for multiple comparisons) in the multiple linear regression analyses. Therefore, it may be possible that only few areas decisively convey the connection between nodal efficiency/local clustering and general intelligence.

There are other possible reasons for the lack of replicable associations between graph metrics and *g* among data sets. First, the samples used for our study vary in multiple demographic details, most importantly age span. While the RUB and NKI samples cover wide age ranges, the HCP and UMN samples comprise young to middle aged adults. Previous studies indicated that resting-state connectivity changes with age and that age is associated with decreasing segregation of brain systems ^[76, 77]^. We regressed age, along with sex and its age-interactions, from our data for analysis, but, when associations are reciprocal, this can remove relevant as well as irrelevant variance. In addition, the samples vary in other demographic factors such as sex ratio, handedness proportion, ethnic composition, and likely most importantly, average IQ scores. The latter in particular may have reduced measurable associations due to range restriction, though range restriction can also enhance them ^[78]^. As three of our four samples show above average intelligence, our results might not be applicable to groups with average or low intelligence. However, many other studies in this area are also characterized by samples with similarly above average mean IQ scores ^[21, 36, 37]^.

Second, the data sets vary in temporal resolution of functional imaging. Even though related images were pre-processed and analyzed in the same manner, this could make for crucial differences. Network neuroscience theory ^[79]^ suggests that intelligent thinking relies on the flexibility of functional networks and ability to switch among various network states quickly. To capture this flexibility, measurements with high temporal resolution are needed. Our study cannot make inferences about flexibility, as we analyzed the complete resting-state sequence instead of conducting time-based analyses. However, it is interesting that data sets with higher temporal resolution (HCP and NKI) revealed more brain areas that were associated with *g* and exhibited greater overlap between results than those with lower temporal resolution (RUB and UMN). Thus, future studies should employ functional imaging with high temporal resolution and examine the flexibility of resting-state networks.

Third, the data sets vary in size. The relations between resting-state connectivity and *g* might be small and some or even all our samples underpowered to detect replicable results. Considerably larger data sets or different approaches might be necessary to reveal such subtle relations even though their impact on intelligence might be limited. One promising approach for future studies would be to investigate task-based functional connectivity networks, as their associations with behavior are usually stronger than those of resting-state networks ^[80, 81]^ Moreover, task-based approaches may provide more insight into the mechanisms behind network states and reveal which specific tasks or cognitive domains are dependent on functional flexibility. Another approach to refine analyses is to employ individualized localization methods, which are based on subject-specific resting-state measurements ^[82]^ In more detail, these methods utilize individual resting-state patterns to individualize brain atlases, which are typically averaged in their discovery cohorts. Examples of this kind of parcellation are multi-session hierarchical Bayesian modeling (MS_HBM) ^[83]^ and group prior individual parcellation (GPIP) ^[84]^. When applied, MS_HBM tends to increase associations between resting-state functional connectivity and cognitive performance. Similar results can be obtained via techniques such as hyper-alignment, which identifies voxels with a similar pattern of neural activity and defines them as neural units ^[85]^. While these methods have been applied to the HCPMMP atlas used in this study, we refrained from using resting state-based individual parcellation for several reasons. First, our aim was to identify causes of inconsistent results in previous studies. Thus, we wanted to maximize comparability among our samples. Second, HCPMMP is a multi-modal brain atlas and its construction did not rely on resting state data alone. Thus, HCPMMP would require a multi-modal individual parcellation. Furthermore, while individualized localization increases effect sizes, it is not clear whether it also increases replicability. Nevertheless, we highly encourage future studies to apply individualized localization to resting state-based brain atlases such as e.g. Schaefer et al. ^[75]^.

In conclusion, our results indicated that graph-theoretical measures of resting-state functional connectivity do not reliably associate with *g*, at least not in specific brain regions. While we do hypothesize that functional connectivity is associated with intelligence, other methods are needed to articulate these relations, if they play a role in intelligent thinking. We suggest using time series-based approaches and investigating the flexibility of functional brain networks via imaging with high temporal resolution. Furthermore, we encourage future studies to focus more on task-based functional connectivity instead of resting-state functional connectivity, and to apply individualized localization methods.

## 4. Data availability

The data from Ruhr University Bochum (RUB) and University of Minnesota (UMN) underlying this study are available from the corresponding author upon reasonable request. The Human Connectome Project (HCP) sample is part of the S1200 release provided by the HCP and can be accessed via its ConnectomeDB platform (https://db.humanconnectome.org). The Nathan Kline Institute (NKI) sample is part of the NKI Rockland Sample release and can be accessed via (http://fcon_1000.projects.nitrc.org/indi/enhanced/).

## 6 Acknowledgements

The authors thank all research assistants for their support during the behavioral measurements. Further, the authors thank PHILIPS Germany (Burkhard Mädler) for the scientific support with the MRI measurements as well as Tobias Otto for technical assistance.

## 7. Author contributions

E.G. conceived the project and supervised the experiments. E.G., O.G., C.F., C.Sc., D.M., C.St., W.J., and C.G.D. designed the project. D.M. and C.St. analyzed the data and wrote the manuscript text. C.F., C.Sc., and C.G.D. collected the data. W.J. calculated the *g* factor scores. All authors edited the manuscript.

## 8. Additional information

This work was supported by the Deutsche Forschungsgemeinschaft (GU 227/16-1). The authors declare no competing interests.

