## Supplementary material for "Investigating robust associations between functional connectivity based on graph theory and general intelligence": Supplementary_Material.docx

1. **Intelligence measurement**

A detailed description of all intelligence measures used for calculation of *g* can be found in Stammen et al. ^[1]^. The following section will give a short overview of all used tests.

1.1. ***RUB Sample***

Intelligence was assessed using the Intelligenz-Struktur-Test 2000 R (I-S-T 2000 R) ^[2]^, a well-established German intelligence test battery ^[2,3]^ that is largely comparable to the Wechsler Adult Intelligence Scale (WAIS) ^[4]^, the Bochumer Matrizentest (BOMAT) ^[5]^, a non-verbal German intelligence test that is comparable to the internationally established Raven’s Advanced Progressive Matrices ^[6]^, the Bochumer Wissenstest (BOWIT) ^[7]^, a German questionnaire assessing general knowledge, and the Zahlenverbindungstest (ZVT) ^[8]^, a trail making test that measures cognitive processing speed.

1.2. ***HCP Sample***

Intelligence was measured with the four subtests Penn Matrix Reasoning Task (PMAT) assessing reasoning ability, Short Penn Continuous Performance Test (SCPT) assessing visual attention, Variable Short Penn Line Orientation Test (VSPLOT) assessing visual-spatial processing, and Penn Word Memory Test (IWRD) assessing verbal episodic memory from the University of Pennsylvania Computerized Neurocognitive Battery (Penn CNB) ^[9,10]^ as well as the seven subtests Flanker Inhibitory Control and Attention Test (Flanker) assessing executive functions (attention), Dimensional Change Card Sort Test (CardSort) assessing executive functions (cognitive flexibility), List Sorting Working Memory Test (ListSort) assessing working memory capacity, Picture Sequence Memory Test (PicSeq) assessing episodic memory, Oral Reading Recognition Test (ReadEng) assessing reading decoding skills, Picture Vocabulary Test (PicVocab) assessing vocabulary knowledge, and Pattern Comparison Processing Speed Test (ProcSpeed) assessing processing speed from the NIH Toolbox for the Assessment of Neurological and Behavioral Function (http://www.nihtoolbox.org) ^[11,12]^.

1.3. ***UMN Sample***

The UMN sample conducted the subtests Block Design (WAIS_BD) and Matrix Reasoning (WAIS_MR) to assess perceptual reasoning, Similarities (WAIS_SIM) and Vocabulary (WAIS_VC) to assess verbal comprehension, and Coding (WAIS_CD) to assess processing speed from the fourth edition Wechsler Adult Intelligence Scale (WAIS-IV) ^[13]^.

1.4. ***NKI Sample***

Intelligence was measured using the four subtests Block Design (WASI_BD), Matrix Reasoning (WASI_MR), Similarities (WASI_SIM), and Vocabulary (WASI_VC) of the second edition Wechsler Abbreviated Scale of Intelligence (WASI-II) ^[14]^, which are conceptually comparable to the subtests from the WAIS-IV (see UMN Sample). However, WASI-II uses unique test items ^[15]^.

2. **References**

1	Stammen, C. *et al.* Robust associations between white matter microstructure and general intelligence. *Cereb Cortex.* **33**, 6723-6741 (2023).

2	Liepmann, D., Beauducel, A., Brocke, B. & Amthauer, R. Intelligenz-Struktur-Test 2000 R (I-S-T 2000 R). Manual., Vol. 2., erweiterte und überarbeitete Auflage (Hogrefe, 2007).

3	Beauducel, A., Brocke, B. & Liepmann, D. Perspectives on fluid and crystallized intelligence: facets for verbal, numerical, and figural intelligence. *Pers. Individ. Differ.* **30**, 977-994 (2001).

4	Erdodi, L. A. *et al.* Wechsler Adult Intelligence Scale-Fourth Edition (WAIS-IV) processing speed scores as measures of noncredible responding: The third generation of embedded performance validity indicators. *Psychol Assess.* **29**, 148-157 (2017).

5	Hossiep, R., Hasella, M. & Turck, D. BOMAT-advanced-short version: Bochumer Matrizentest. (Hogrefe, 2001).

6	Raven, J. C., Court, J. H. & Raven, J. Coloured progressive matrices. Manual for Raven’s Progressive Matrices and Vocabulary Scales. (Oxford Psychologists Press, 1990).

7	Hossiep, R. & Schulte, M. BOWIT: Bochumer Wissenstest. (Hogrefe, 2008).

8	Oswald, W. D. & Roth, E. Der Zahlen-Verbindungs-Test (ZVT). (Hogrefe Verlag für Psychologie, 1987).

9	Gur, R. C. *et al.* Computerized neurocognitive scanning: I. Methodology and validation in healthy people. *NPP.* **25**, 766-776 (2001).

10	Gur, R. C. *et al.* A cognitive neuroscience-based computerized battery for efficient measurement of individual differences: standardization and initial construct validation. *J Neurosci Methods.* **187**, 254-262 (2010).

11	Gershon, R. C. *et al.* NIH Toolbox for the assessment of neurological and behavioral function. *Neurology.* **80**, S2-S6 (2013).

12	Weintraub, S. *et al.* Cognition assessment using the NIH Toolbox. *Neurology.* **80**, S54-S64 (2013).

13	Wechsler, D. Wechsler Adult Intelligence Scale - Forth edition (WAIS-IV). (Pearson Assessment, 2008).

14	Wechsler, D. Wechsler Abbreviated Intelligence Scale - Second edition (WASI-II). (NCS Pearson, 2011).

15	McCrimmon, A. W. & Smith, A. D. Review of the Wechsler Abbreviated Scale of Intelligence, Second Edition (WASI-II). *J Psychoeduc Assess.* **31**, 337-341 (2012).
