## Supplementary material for "Investigating robust associations between functional connectivity based on graph theory and general intelligence": Supplementary_Tables.pdf

**Supplementary Table S1** List of all HCPMMP and subcortical areas with non-zero effect sizes for each data set (nodal efficiency). Left hemispheric areas are labeled "L", right hemispheric areas are labled "R". HCP sessions are depicted in blue, NKI sessions in orange, UMN session in yellow and RUB session in green.

| HCP day 1 | | HCP day 2 | | HCP day 1 session 1 | | HCP day 1 session 2 | | HCP day 1 & 2 | | NKI (TR = 645) | | NKI (TR = 1400) | | NKI (TR = 2500) | | UMN | |
|---|---|---|---|---|---|---|---|---|---|---|---|---|---|---|---|---|---|
| **Area** | **beta** | **Area** | **beta** | **Area** | **beta** | **Area** | **beta** | **Area** | **beta** | **Area** | **beta** | **Area** | **beta** | **Area** | **beta** | **Area** | **beta** |
| L_V6 | -0,008 | L_SFL | -0,014 | L_V6 | -0,033 | L_MST | 0,005 | L_PSL | -0,065 | L_3b | -0,023 | L_MST | 0,021 | L_FEF | -0,007 | L_V4 | -0,097 |
| L_RSC | -0,037 | L_6ma | -0,056 | L_MT | -0,002 | L_V8 | -0,027 | L_23d | 0,001 | L_FEF | -0,003 | L_POS2 | -0,027 | L_PEF | -0,004 | L_STV | -0,001 |
| L_MT | -0,040 | L_6d | -0,048 | L_A1 | -0,020 | L_FFC | -0,057 | L_24dd | -0,006 | L_5m | -0,047 | L_p24pr | 0,000 | L_LO2 | -0,092 | L_31pv | 0,034 |
| L_SFL | -0,052 | L_6v | 0,000 | L_SFL | -0,045 | L_PCV | -0,048 | L_1 | -0,001 | L_5mv | -0,109 | L_6r | 0,189 | L_PCV | -0,053 | L_LIPv | -0,030 |
| L_PCV | -0,029 | L_p24pr | 0,001 | L_PCV | -0,024 | L_7m | 0,065 | L_p32pr | 0,031 | L_10r | 0,013 | L_EC | -0,047 | L_7Pm | -0,005 | L_IFJp | -0,109 |
| L_7m | 0,055 | L_a24pr | 0,002 | L_7m | 0,011 | L_23d | 0,025 | L_6r | 0,021 | L_8BL | 0,019 | L_PreS | -0,043 | L_5mv | -0,058 | L_13l | 0,005 |
| L_23d | 0,041 | L_p32pr | 0,029 | L_7PL | 0,000 | L_v23ab | -0,014 | L_a10p | 0,061 | L_6r | 0,031 | L_H | -0,037 | L_5L | 0,000 | L_47s | 0,086 |
| L_5L | -0,023 | L_8BM | 0,011 | L_6v | -0,036 | L_SCEF | -0,039 | L_11l | 0,032 | L_10v | 0,051 | L_PeEc | -0,004 | L_6r | 0,074 | L_FOP3 | 0,061 |
| L_SCEF | -0,007 | L_a47r | 0,052 | L_33pr | -0,044 | L_7PC | -0,015 | L_47s | 0,064 | L_LIPd | -0,003 | L_STGa | -0,062 | L_9.46d | -0,073 | L_AIP | -0,001 |
| L_7PC | -0,077 | L_IFJp | 0,017 | L_a24 | -0,026 | L_6mp | -0,001 | L_i6.8 | -0,006 | L_EC | -0,072 | L_TF | -0,023 | L_LIPd | -0,008 | L_STSdp | -0,044 |
| L_33pr | -0,039 | L_IFSp | 0,031 | L_p32 | -0,017 | L_a24 | -0,002 | L_RI | -0,010 | L_H | -0,008 | L_STSva | 0,085 | L_OP2.3 | -0,062 | L_TF | 0,056 |
| L_a24 | -0,019 | L_IFSa | 0,056 | L_47m | 0,053 | L_a47r | 0,020 | L_MI | -0,005 | L_PeEc | -0,004 | R_V1 | -0,007 | L_PFcm | 0,023 | L_PGp | 0,008 |
| L_p32 | -0,019 | L_a9.46v | 0,053 | L_45 | -0,015 | L_IFJp | 0,030 | L_AVI | -0,074 | L_DVT | -0,010 | R_POS2 | -0,017 | L_PoI2 | 0,016 | L_PF | -0,059 |
| L_47m | 0,035 | L_a10p | 0,005 | L_a47r | 0,049 | L_46 | 0,005 | L_PreS | -0,025 | L_PGi | 0,006 | R_LO1 | 0,014 | L_PFt | -0,001 | L_25 | 0,028 |
| L_47l | 0,015 | L_LIPd | 0,011 | L_IFJa | 0,061 | L_13l | 0,088 | L_STSda | 0,019 | L_pOFC | -0,038 | R_31pv | 0,049 | L_EC | -0,041 | L_PI | -0,028 |
| L_6r | 0,043 | L_PoI2 | -0,037 | L_IFJp | 0,010 | L_47s | 0,021 | L_TE2a | 0,000 | L_p47r | 0,077 | R_5mv | -0,182 | L_DVT | -0,016 | R_V4 | -0,120 |
| L_IFJa | 0,001 | L_AVI | -0,009 | L_a10p | 0,015 | L_LIPd | 0,008 | L_s32 | 0,031 | L_A4 | 0,038 | R_47l | 0,017 | L_s32 | -0,049 | R_SFL | 0,031 |
| L_IFJp | 0,060 | L_PreS | -0,051 | L_47s | 0,052 | L_MI | -0,027 | L_p10p | 0,016 | R_V6 | -0,042 | R_a47r | -0,002 | L_Ig | -0,041 | R_2 | -0,008 |
| L_9.46d | -0,016 | L_TPOJ3 | 0,004 | L_AAIC | -0,011 | L_PreS | -0,081 | R_LO2 | 0,011 | R_24dd | -0,003 | R_IFJp | 0,085 | L_A4 | 0,064 | R_IFJp | 0,063 |
| L_a10p | 0,011 | L_PHA2 | 0,023 | L_PHA3 | -0,018 | L_STSvp | 0,066 | R_A1 | -0,008 | R_6ma | 0,004 | R_9a | -0,017 | L_TE1m | 0,101 | R_46 | 0,040 |
| L_10pp | -0,037 | L_A4 | -0,005 | L_TE2p | -0,013 | L_TPOJ3 | 0,070 | R_5mv | -0,008 | R_6d | -0,057 | R_PHA1 | -0,100 | L_a32pr | -0,001 | R_52 | 0,041 |
| L_11l | 0,004 | L_PI | -0,001 | L_TPOJ3 | 0,013 | L_IP0 | 0,023 | R_5L | 0,016 | R_6v | 0,011 | R_STSvp | 0,016 | R_V8 | 0,002 | R_MI | 0,059 |
| L_OFC | -0,041 | R_55b | -0,049 | L_PGs | 0,020 | L_PGs | 0,003 | R_a24pr | -0,029 | R_8BL | 0,081 | R_TE2a | -0,011 | R_FFC | 0,062 | R_AAIC | 0,008 |
| L_47s | 0,027 | R_7m | 0,026 | L_VMV1 | -0,015 | L_V3CD | 0,011 | R_6r | 0,026 | R_10d | 0,034 | R_PFop | 0,050 | R_A1 | -0,002 | R_H | -0,043 |
| L_LIPd | 0,016 | R_33pr | -0,040 | L_LO3 | -0,002 | L_31pd | 0,054 | R_IFJp | 0,001 | R_44 | 0,016 | R_V6A | 0,010 | R_v23ab | -0,010 | R_PBelt | -0,081 |
| L_6a | 0,008 | R_IFSp | -0,011 | L_VMV2 | -0,030 | L_VVC | -0,028 | R_IFSp | -0,045 | R_H | -0,063 | R_Ig | -0,008 | R_5mv | -0,067 | R_STSvp | -0,040 |
| L_RI | -0,041 | R_OFC | -0,005 | L_31pd | 0,018 | L_25 | -0,086 | R_IFSa | -0,024 | R_TE2a | -0,051 | R_p24 | 0,004 | R_7PL | 0,017 | R_PF | 0,016 |

| | | | | | | | | | | | | | | | | | |
|---|---|---|---|---|---|---|---|---|---|---|---|---|---|---|---|---|---|
| L_MI | -0,080 | R_MI | -0,003 | L_31a | 0,040 | R_V6 | 0,054 | R_11l | 0,009 | R_31a | -0,033 | L_accumbens | -0,070 | R_44 | 0,059 | R_TE1m | -0,018 |
| L_AVI | -0,015 | R_PHA3 | -0,023 | L_VVC | -0,048 | R_FEF | -0,030 | R_13l | -0,023 | | | L_hippo | -0,010 | R_45 | 0,021 | R_p24 | 0,102 |
| L_FOP1 | -0,027 | R_TF | -0,003 | L_s32 | -0,077 | R_55b | -0,008 | R_s6.8 | 0,072 | | | R_accumbens | -0,023 | R_IFSp | 0,114 | L_accumbens | 0,019 |
| L_PreS | -0,042 | R_IP2 | 0,035 | L_FOP5 | -0,013 | R_V7 | 0,037 | R_OP1 | -0,006 | | | R_putamen | -0,037 | R_52 | 0,052 | L_hippocampus | -0,010 |
| L_STGa | 0,024 | L_putamen | -0,014 | L_TE1m | 0,070 | R_V3B | 0,033 | R_TA2 | 0,000 | | | | | R_FOP3 | 0,060 | | |
| L_STSda | 0,008 | R_putamen | -0,023 | L_a32pr | -0,050 | R_23d | 0,047 | R_PHA1 | -0,041 | | | | | R_EC | -0,078 | | |
| L_STSvp | 0,052 | | | L_p24 | -0,036 | R_6v | -0,027 | R_PHA3 | -0,006 | | | | | R_PreS | -0,102 | | |
| L_TE1p | -0,011 | | | R_V6 | 0,129 | R_a24 | -0,042 | R_TGd | 0,010 | | | | | R_PeEc | -0,008 | | |
| L_TF | 0,014 | | | R_PEF | 0,008 | R_p32 | -0,026 | R_TE2a | 0,002 | | | | | R_TE1a | 0,019 | | |
| L_TE2p | -0,024 | | | R_RSC | -0,001 | R_10r | -0,015 | R_PF | -0,016 | | | | | R_TE1p | -0,132 | | |
| L_TPOJ3 | 0,048 | | | R_LO2 | -0,038 | R_IFJp | 0,009 | R_VMV1 | 0,006 | | | | | R_PFop | 0,109 | | |
| L_PGp | -0,010 | | | R_A1 | -0,004 | R_10v | 0,015 | R_FOP5 | -0,034 | | | | | R_VVC | 0,022 | | |
| L_IP2 | 0,002 | | | R_POS1 | -0,001 | R_OFC | 0,001 | L_caudat | 0,014 | | | | | R_pOFC | -0,012 | | |
| L_PFop | 0,093 | | | R_7Am | -0,022 | R_OP1 | 0,034 | | | | | | | R_Ig | -0,066 | | |
| L_PF | 0,004 | | | R_VIP | -0,011 | R_TF | -0,013 | | | | | | | R_TE1m | 0,013 | | |
| L_PFm | 0,029 | | | R_33pr | -0,016 | R_TPOJ3 | 0,015 | | | | | | | R_accum | -0,004 | | |
| L_LO3 | -0,035 | | | R_10v | 0,062 | R_IP2 | 0,052 | | | | | | | R_amygd | 0,017 | | |
| L_VVC | -0,062 | | | R_13l | 0,103 | R_VMV3 | -0,041 | | | | | | | | | | |
| L_PoI1 | 0,004 | | | R_OP1 | 0,047 | R_31pd | 0,002 | | | | | | | | | | |
| L_p10p | 0,026 | | | R_FOP4 | 0,011 | R_25 | -0,018 | | | | | | | | | | |
| L_PI | -0,004 | | | R_Pir | -0,021 | R_pOFC | -0,026 | | | | | | | | | | |
| L_a32pr | 0,065 | | | R_STGa | 0,020 | R_TE1m | 0,010 | | | | | | | | | | |
| R_V6 | 0,071 | | | R_TE1a | 0,003 | L_accumbens | -0,055 | | | | | | | | | | |
| R_FEF | -0,014 | | | R_IP2 | 0,049 | L_ventraldc | 0,007 | | | | | | | | | | |
| R_V3B | 0,009 | | | R_25 | -0,072 | R_accumbens | -0,018 | | | | | | | | | | |
| R_SFL | 0,022 | | | R_s32 | -0,022 | R_caudate | 0,000 | | | | | | | | | | |
| R_PCV | -0,009 | | | R_PI | -0,004 | | | | | | | | | | | | |
| R_7Pm | -0,015 | | | L_accumben | -0,048 | | | | | | | | | | | | |
| R_31pv | 0,046 | | | L_ventraldc | 0,003 | | | | | | | | | | | | |
| R_23c | -0,029 | | | | | | | | | | | | | | | | |
| R_7PC | 0,014 | | | | | | | | | | | | | | | | |
| R_LIPv | 0,018 | | | | | | | | | | | | | | | | |
| R_6v | -0,016 | | | | | | | | | | | | | | | | |
| R_33pr | -0,016 | | | | | | | | | | | | | | | | |

| | |
|---|---|
| R_a24pr | -0,004 |
| R_10r | -0,013 |
| R_8Av | 0,038 |
| R_9p | -0,001 |
| R_IFSp | -0,020 |
| R_13l | -0,010 |
| R_47s | 0,044 |
| R_i6.8 | 0,007 |
| R_43 | -0,054 |
| R_PFcm | -0,019 |
| R_TA2 | 0,005 |
| R_FOP4 | 0,030 |
| R_MI | -0,047 |
| R_STGa | 0,041 |
| R_A5 | 0,017 |
| R_PHA3 | -0,012 |
| R_TE2a | 0,010 |
| R_TF | -0,022 |
| R_IP2 | 0,051 |
| R_VMV1 | 0,074 |
| R_VMV3 | -0,081 |
| R_A4 | 0,003 |
| R_PI | -0,011 |
| L_hippo | 0,018 |
| L_pallidu | -0,028 |
| L_putam | -0,015 |
| L_ventral | 0,031 |
| R_caudat | -0,002 |
| R_putam | -0,001 |
| R_ventra | -0,040 |

**Supplementary Table S2** List of all HCPMMP and subcortical areas overlapping between different datasets (nodal efficiency, see Figure 2). Left hemispheric areas are labeled "L", right hemispheric areas are labled "R". Beta refers to the mean beta of the overlapping datasets.

| | | | | | | | | | |
|---|---|---|---|---|---|---|---|---|---|
| **HCP across days** | | **HCP same day** | | **NKI** | | **TR ≤ 720 ms data sets** | | **TR = 2000 ms data sets** | |
| **Area** | **beta** | **Area** | **beta** | **Area** | **beta** | **Area** | **beta** | **Area** | **beta** |
| L_SFL | -0,033 | L_PCV | -0,036 | L_6r | 0,098 | L_6r | 0,0259 | - | - |
| L_IFJp | 0,039 | L_7m | 0,038 | L_EC | -0,053 | R_TE2a | -0,0246 | | |
| L_a10p | 0,008 | L_a24 | -0,014 | | | | | | |
| L_LIPd | 0,014 | L_a47r | 0,034 | | | | | | |
| L_AVI | -0,012 | L_IFJp | 0,020 | | | | | | |
| L_PreS | -0,047 | L_47s | 0,037 | | | | | | |
| L_TPOJ3 | 0,026 | L_TPOJ3 | 0,041 | | | | | | |
| L_PI | -0,003 | L_PGs | 0,011 | | | | | | |
| R_33pr | -0,028 | L_31pd | 0,036 | | | | | | |
| R_IFSp | -0,015 | L_VVC | -0,038 | | | | | | |
| R_MI | -0,025 | R_V6 | 0,092 | | | | | | |
| R_PHA3 | -0,017 | R_10v | 0,038 | | | | | | |
| R_TF | -0,012 | R_OP1 | 0,041 | | | | | | |
| R_IP2 | 0,043 | R_IP2 | 0,050 | | | | | | |
| L_putamen | -0,014 | R_25 | -0,045 | | | | | | |
| R_putamen | -0,012 | L_accumbens | -0,052 | | | | | | |
| | | L_ventraldc | 0,005 | | | | | | |

**Supplementary Table S3** List of all HCPMMP and subcortical areas with non-zero effect sizes for each data set (local clustering). Left hemispheric areas are labeled "L", right hemispheric areas are labled "R". HCP sessions are depicted in blue, NKI sessions in orange, UMN session in green and RUB session in green.

| HCP day 1 | | HCP day 2 | | CP day 1 session | | HCP day 1 session 2 | | HCP day 1 & 2 | | NKI (TR = 645) | | NKI (TR = 1400) | | NKI (TR = 2500) | | UMN | |
|---|---|---|---|---|---|---|---|---|---|---|---|---|---|---|---|---|---|
| Area | beta | Area | beta | Area | beta | Area | beta | Area | beta | Area | beta | Area | beta | Area | beta | Area | beta |
| L_V1 | -0,005 | L_MST | -0,018 | L_V8 | -0,009 | L_8BM | 0,038 | L_V2 | 0,015 | L_POS2 | -0,040 | L_SFL | 0,016 | L_V3 | 0,015 | L_7PC | -0,029 |
| L_MST | -0,006 | L_V6 | 0,002 | L_POS2 | -0,016 | L_8BL | 0,006 | L_55b | -0,032 | L_SFL | 0,047 | L_23d | 0,041 | L_PI | -0,002 | L_9a | 0,015 |
| L_SFL | 0,002 | L_SFL | 0,019 | L_FFC | -0,044 | L_45 | 0,009 | L_RSC | -0,005 | L_5L | -0,023 | L_23c | -0,002 | L_a24p | 0,000 | | |
| L_STV | -0,010 | L_d23ab | -0,005 | L_v23ab | 0,034 | L_IFSp | 0,009 | L_LO2 | 0,002 | L_7PC | 0,021 | L_24dd | 0,006 | L_R | 0,010 | | |
| L_v23ab | 0,007 | L_SCEF | -0,013 | L_6d | 0,019 | L_47s | 0,018 | L_A1 | -0,002 | L_3a | 0,005 | L_d32 | -0,019 | L_STSv | 0,014 | | |
| L_5mv | -0,005 | L_7Am | -0,048 | L_6v | 0,029 | L_LIPd | 0,009 | L_SFL | 0,019 | L_47m | 0,008 | L_44 | 0,010 | R_SF | -0,030 | | |
| L_7PC | -0,034 | L_1 | 0,005 | L_33pr | -0,010 | L_STSdp | 0,002 | L_31pv | -0,018 | L_47l | 0,037 | L_a47r | -0,029 | R_3 | 0,010 | | |
| L_6v | 0,017 | L_a24pr | 0,003 | L_8BM | 0,042 | L_TE1p | 0,006 | L_24dd | -0,005 | L_6a | 0,006 | L_OFC | 0,001 | R_d3 | -0,068 | | |
| L_8BM | 0,023 | L_8BM | 0,033 | L_47m | 0,036 | L_TPOJ1 | 0,021 | L_LIPv | 0,008 | L_43 | 0,017 | L_LIPd | 0,007 | R_10 | 0,000 | | |
| L_45 | 0,052 | L_8BL | 0,005 | L_8BL | 0,010 | L_25 | -0,019 | L_1 | 0,002 | L_PFt | 0,030 | L_6a | 0,000 | R_6 | 0,033 | | |
| L_IFSa | 0,000 | L_44 | 0,018 | L_a47r | 0,007 | L_TE1m | 0,003 | L_8C | -0,001 | L_STSdp | 0,027 | L_MI | 0,002 | R_IFJ | 0,019 | | |
| L_a10p | 0,047 | L_a10p | 0,008 | L_6r | 0,026 | R_V3B | 0,004 | L_43 | -0,039 | L_STSvp | 0,015 | L_PFt | 0,003 | R_9.46 | -0,011 | | |
| L_13l | -0,014 | L_11l | 0,000 | L_11l | -0,013 | R_AVI | 0,009 | L_AVI | 0,002 | L_TE1p | 0,005 | L_AIP | 0,042 | R_PHA | -0,008 | | |
| L_OP4 | 0,025 | L_OP2.3 | 0,006 | L_47s | 0,072 | R_LO3 | 0,004 | L_PreS | -0,025 | L_LO3 | -0,021 | L_PreS | -0,032 | R_TPOJ | -0,064 | | |
| L_RI | -0,022 | L_PFt | 0,036 | L_TPOJ2 | -0,031 | R_pOFC | -0,004 | L_TPOJ2 | -0,034 | L_TE1m | 0,009 | L_TPOJ1 | 0,017 | R_pOF | -0,006 | | |
| L_MI | 0,018 | L_AIP | 0,007 | L_PGp | -0,015 | R_TE1m | 0,057 | L_PFm | -0,010 | R_p32pr | 0,005 | L_DVT | -0,028 | R_putamen | 0,033 | | |
| L_PGi | 0,006 | L_EC | -0,011 | L_PGi | 0,008 | R_accumbens | -0,011 | L_VVC | -0,008 | R_9.46d | -0,005 | L_PGp | -0,040 | | | | |
| L_VMV3 | -0,006 | L_PreS | -0,045 | L_a32pr | -0,038 | | | L_25 | 0,001 | R_13l | 0,001 | L_PFop | 0,045 | | | | |
| L_s32 | 0,014 | L_PFm | -0,005 | R_V1 | -0,024 | | | L_s32 | 0,033 | R_6a | 0,037 | L_TE1m | 0,018 | | | | |
| L_FOP5 | -0,008 | L_PHA2 | 0,003 | R_V6 | -0,021 | | | L_p47r | -0,051 | R_FOP3 | 0,040 | R_PEF | 0,073 | | | | |
| L_p10p | -0,001 | L_FST | -0,004 | R_V3 | 0,035 | | | L_MBelt | 0,008 | R_FOP2 | 0,018 | R_a24 | -0,014 | | | | |
| L_A4 | 0,022 | L_VVC | -0,017 | R_3b | 0,046 | | | L_A4 | -0,034 | R_TE1a | 0,022 | R_8BM | -0,024 | | | | |
| L_PI | -0,006 | L_TGv | -0,013 | R_24dd | 0,013 | | | R_V3 | 0,022 | R_PGi | 0,004 | R_8Ad | 0,040 | | | | |
| L_p24 | 0,000 | R_IPS1 | 0,023 | R_7PL | 0,003 | | | R_MT | -0,005 | R_31pd | 0,001 | R_9m | -0,069 | | | | |

| | | | | | | | | | | | |
|---|---|---|---|---|---|---|---|---|---|---|---|
| R_V1 | -0,003 | R_V3B | 0,018 | R_p24pr | -0,017 | R_7Pm | 0,002 | R_s32 | 0,007 | R_IFJp | 0,021 |
| R_V8 | 0,009 | R_MT | -0,005 | R_p32 | -0,013 | R_5L | 0,000 | R_p24 | -0,059 | R_9.46d | -0,073 |
| R_3b | 0,012 | R_7m | 0,019 | R_OFC | 0,019 | R_1 | 0,007 | R_accumbens | -0,001 | R_9a | -0,011 |
| R_v23ab | 0,006 | R_5m | 0,006 | R_FOP4 | 0,018 | R_a24 | 0,015 | | | R_10pp | -0,003 |
| R_5m | 0,036 | R_3a | 0,010 | R_Pir | -0,002 | R_p32 | 0,001 | | | R_52 | 0,030 |
| R_MIP | 0,002 | R_8BL | 0,016 | R_FOP3 | 0,005 | R_45 | -0,017 | | | R_MI | 0,030 |
| R_3a | 0,008 | R_47l | 0,010 | R_FOP2 | 0,008 | R_IFSa | -0,007 | | | R_FOP1 | 0,044 |
| R_8Av | 0,006 | R_11l | -0,042 | R_STSdp | 0,010 | R_46 | 0,020 | | | R_FOP3 | 0,007 |
| R_8BL | 0,001 | R_47s | 0,001 | R_TE2a | 0,023 | R_10v | 0,044 | | | R_AIP | 0,006 |
| R_8C | 0,017 | R_i6.8 | 0,002 | R_IP2 | 0,019 | R_13l | -0,038 | | | R_PreS | -0,008 |
| R_9.46d | -0,004 | R_s6.8 | 0,038 | R_IP0 | 0,023 | R_6a | 0,020 | | | R_TE2p | 0,003 |
| R_10v | 0,000 | R_AAIC | -0,024 | R_PFm | 0,000 | R_43 | 0,017 | | | R_PHT | -0,018 |
| R_13l | -0,024 | R_PFt | -0,002 | R_PGi | 0,006 | R_FOP2 | 0,034 | | | R_31pd | 0,025 |
| R_FOP4 | 0,001 | R_STSvp | 0,006 | R_PGs | 0,000 | R_PBelt | 0,023 | | | R_VVC | 0,057 |
| R_Pir | 0,016 | R_TE1a | 0,052 | R_V3CD | 0,013 | R_A5 | -0,009 | | | R_p10p | -0,026 |
| R_AVI | 0,010 | R_PFm | 0,000 | R_s32 | -0,010 | R_PHA3 | -0,041 | | | | |
| R_EC | -0,009 | R_V3CD | 0,042 | R_TGv | 0,007 | R_STSda | -0,015 | | | | |
| R_TE1a | 0,035 | R_25 | 0,021 | | | R_TE1a | 0,010 | | | | |
| R_TE2a | -0,007 | R_p10p | -0,006 | | | R_TPOJ2 | -0,039 | | | | |
| R_PF | -0,004 | R_TGv | -0,014 | | | R_V4t | 0,031 | | | | |
| R_PGs | 0,027 | R_PI | -0,012 | | | R_25 | 0,015 | | | | |
| R_VMV1 | 0,010 | L_hippo | 0,016 | | | R_pOFC | -0,023 | | | | |
| R_VMV3 | -0,032 | L_putamen | -0,022 | | | R_FOP5 | -0,024 | | | | |
| R_V3CD | 0,037 | R_hippocampus | 0,024 | | | L_pallidum | 0,007 | | | | |
| R_FOP5 | -0,042 | | | | | L_thalamus_prop | 0,004 | | | | |
| R_a32pr | -0,002 | | | | | R_amygdala | 0,038 | | | | |
| R_caudate | -0,026 | | | | | | | | | | |
| R_ventraldc | -0,042 | | | | | | | | | | |

**Supplementary Table S4** List of all HCPMMP and subcortical areas overlapping between different datasets (local efficiency, see Figure 3). Left hemispheric areas are labeled "L", right hemispheric areas are labled "R".

| HCP across days | | HCP same day | | NKI | | TR ≤ 720 ms data sets | | TR = 2000 ms data sets | |
|---|---|---|---|---|---|---|---|---|---|
| **Area** | **beta** | **Area** | **beta** | **Area** | **beta** | **Area** | **beta** | **Area** | **beta** |
| L_MST | -0,012 | L_8BM | 0,040 | L_6r | 0,098 | L_SFL | 0,033 | - | - |
| L_SFL | 0,011 | L_8BL | 0,008 | | | L_43 | -0,011 | | |
| L_8BM | 0,028 | L_47s | 0,045 | | | R_13l | -0,019 | | |
| L_a10p | 0,027 | | | | | R_6a | 0,029 | | |
| R_5m | 0,021 | | | | | R_FOP2 | 0,026 | | |
| R_3a | 0,009 | | | | | R_TE1a | 0,016 | | |
| R_8BL | 0,009 | | | | | | | | |
| R_TE1a | 0,043 | | | | | | | | |
| R_V3CD | 0,040 | | | | | | | | |

**Supplementary Table S5** Reliability in terms of ICC of all HCPMMP and subcortical areas (nodal efficiency & local clustering). Left hemispheric areas are labeled "L", right hemispheric areas are labeled "R".

| Nodal Efficiency | | | | | | | Local Clustering | | | | | |
|---|---|---|---|---|---|---|---|---|---|---|---|---|
| HCP across days | | HCP same day | | NKI | | | HCP across days | | HCP same day | | NKI | |
| **Area** | **ICC** | **Area** | **ICC** | **Area** | **ICC** | | **Area** | **ICC** | **Area** | **ICC** | **Area** | **ICC** |
| L_V1 | 0,431 | L_V1 | 0,561 | L_V1 | 0,407 | | L_V1 | 0,376 | L_V1 | 0,446 | L_V1 | 0,355 |
| L_MST | 0,576 | L_MST | 0,749 | L_MST | 0,470 | | L_MST | 0,324 | L_MST | 0,454 | L_MST | 0,330 |
| L_V6 | 0,513 | L_V6 | 0,679 | L_V6 | 0,453 | | L_V6 | 0,428 | L_V6 | 0,406 | L_V6 | 0,257 |
| L_V2 | 0,474 | L_V2 | 0,605 | L_V2 | 0,441 | | L_V2 | 0,367 | L_V2 | 0,392 | L_V2 | 0,324 |
| L_V3 | 0,449 | L_V3 | 0,607 | L_V3 | 0,391 | | L_V3 | 0,424 | L_V3 | 0,382 | L_V3 | 0,431 |
| L_V4 | 0,436 | L_V4 | 0,609 | L_V4 | 0,434 | | L_V4 | 0,467 | L_V4 | 0,440 | L_V4 | 0,358 |
| L_V8 | 0,427 | L_V8 | 0,639 | L_V8 | 0,411 | | L_V8 | 0,353 | L_V8 | 0,322 | L_V8 | 0,327 |
| L_4 | 0,586 | L_4 | 0,677 | L_4 | 0,534 | | L_4 | 0,346 | L_4 | 0,326 | L_4 | 0,331 |
| L_3b | 0,564 | L_3b | 0,666 | L_3b | 0,483 | | L_3b | 0,373 | L_3b | 0,384 | L_3b | 0,249 |
| L_FEF | 0,500 | L_FEF | 0,644 | L_FEF | 0,520 | | L_FEF | 0,227 | L_FEF | 0,304 | L_FEF | 0,165 |
| L_PEF | 0,554 | L_PEF | 0,713 | L_PEF | 0,565 | | L_PEF | 0,295 | L_PEF | 0,298 | L_PEF | 0,157 |
| L_55b | 0,545 | L_55b | 0,655 | L_55b | 0,526 | | L_55b | 0,315 | L_55b | 0,320 | L_55b | 0,207 |
| L_V3A | 0,481 | L_V3A | 0,627 | L_V3A | 0,451 | | L_V3A | 0,468 | L_V3A | 0,479 | L_V3A | 0,388 |
| L_RSC | 0,412 | L_RSC | 0,542 | L_RSC | 0,494 | | L_RSC | 0,246 | L_RSC | 0,236 | L_RSC | 0,289 |
| L_POS2 | 0,430 | L_POS2 | 0,629 | L_POS2 | 0,502 | | L_POS2 | 0,334 | L_POS2 | 0,255 | L_POS2 | 0,240 |
| L_V7 | 0,471 | L_V7 | 0,630 | L_V7 | 0,575 | | L_V7 | 0,428 | L_V7 | 0,357 | L_V7 | 0,243 |
| L_IPS1 | 0,491 | L_IPS1 | 0,636 | L_IPS1 | 0,522 | | L_IPS1 | 0,285 | L_IPS1 | 0,360 | L_IPS1 | 0,282 |
| L_FFC | 0,435 | L_FFC | 0,569 | L_FFC | 0,451 | | L_FFC | 0,261 | L_FFC | 0,233 | L_FFC | 0,249 |
| L_V3B | 0,517 | L_V3B | 0,676 | L_V3B | 0,468 | | L_V3B | 0,356 | L_V3B | 0,368 | L_V3B | 0,276 |
| L_LO1 | 0,461 | L_LO1 | 0,645 | L_LO1 | 0,444 | | L_LO1 | 0,445 | L_LO1 | 0,426 | L_LO1 | 0,314 |
| L_LO2 | 0,415 | L_LO2 | 0,601 | L_LO2 | 0,385 | | L_LO2 | 0,389 | L_LO2 | 0,370 | L_LO2 | 0,298 |
| L_PIT | 0,450 | L_PIT | 0,608 | L_PIT | 0,370 | | L_PIT | 0,317 | L_PIT | 0,311 | L_PIT | 0,221 |
| L_MT | 0,497 | L_MT | 0,698 | L_MT | 0,470 | | L_MT | 0,252 | L_MT | 0,435 | L_MT | 0,289 |
| L_A1 | 0,506 | L_A1 | 0,640 | L_A1 | 0,463 | | L_A1 | 0,199 | L_A1 | 0,374 | L_A1 | 0,268 |

| | | | | | | | | | | | | |
|---|---|---|---|---|---|---|---|---|---|---|---|---|
| L_PSL | 0,543 | L_PSL | 0,673 | L_PSL | 0,520 | | L_PSL | 0,274 | L_PSL | 0,389 | L_PSL | 0,250 |
| L_SFL | 0,459 | L_SFL | 0,666 | L_SFL | 0,505 | | L_SFL | 0,395 | L_SFL | 0,359 | L_SFL | 0,070 |
| L_PCV | 0,409 | L_PCV | 0,617 | L_PCV | 0,526 | | L_PCV | 0,229 | L_PCV | 0,291 | L_PCV | 0,205 |
| L_STV | 0,536 | L_STV | 0,678 | L_STV | 0,501 | | L_STV | 0,242 | L_STV | 0,264 | L_STV | 0,291 |
| L_7Pm | 0,488 | L_7Pm | 0,622 | L_7Pm | 0,475 | | L_7Pm | 0,313 | L_7Pm | 0,277 | L_7Pm | 0,170 |
| L_7m | 0,443 | L_7m | 0,644 | L_7m | 0,446 | | L_7m | 0,392 | L_7m | 0,369 | L_7m | 0,317 |
| L_POS1 | 0,417 | L_POS1 | 0,600 | L_POS1 | 0,396 | | L_POS1 | 0,207 | L_POS1 | 0,247 | L_POS1 | 0,246 |
| L_23d | 0,415 | L_23d | 0,497 | L_23d | 0,491 | | L_23d | 0,257 | L_23d | 0,133 | L_23d | 0,257 |
| L_v23ab | 0,432 | L_v23ab | 0,651 | L_v23ab | 0,499 | | L_v23ab | 0,211 | L_v23ab | 0,283 | L_v23ab | 0,328 |
| L_d23ab | 0,393 | L_d23ab | 0,605 | L_d23ab | 0,527 | | L_d23ab | 0,306 | L_d23ab | 0,288 | L_d23ab | 0,336 |
| L_31pv | 0,430 | L_31pv | 0,701 | L_31pv | 0,479 | | L_31pv | 0,355 | L_31pv | 0,347 | L_31pv | 0,297 |
| L_5m | 0,529 | L_5m | 0,660 | L_5m | 0,544 | | L_5m | 0,404 | L_5m | 0,445 | L_5m | 0,319 |
| L_5mv | 0,469 | L_5mv | 0,665 | L_5mv | 0,594 | | L_5mv | 0,300 | L_5mv | 0,272 | L_5mv | 0,197 |
| L_23c | 0,409 | L_23c | 0,631 | L_23c | 0,547 | | L_23c | 0,207 | L_23c | 0,154 | L_23c | 0,090 |
| L_5L | 0,505 | L_5L | 0,633 | L_5L | 0,472 | | L_5L | 0,307 | L_5L | 0,303 | L_5L | 0,226 |
| L_24dd | 0,466 | L_24dd | 0,670 | L_24dd | 0,585 | | L_24dd | 0,249 | L_24dd | 0,299 | L_24dd | 0,291 |
| L_24dv | 0,443 | L_24dv | 0,647 | L_24dv | 0,551 | | L_24dv | 0,228 | L_24dv | 0,326 | L_24dv | 0,197 |
| L_7AL | 0,461 | L_7AL | 0,625 | L_7AL | 0,495 | | L_7AL | 0,316 | L_7AL | 0,339 | L_7AL | 0,235 |
| L_SCEF | 0,434 | L_SCEF | 0,681 | L_SCEF | 0,584 | | L_SCEF | 0,253 | L_SCEF | 0,251 | L_SCEF | 0,156 |
| L_6ma | 0,465 | L_6ma | 0,668 | L_6ma | 0,525 | | L_6ma | 0,234 | L_6ma | 0,266 | L_6ma | 0,149 |
| L_7Am | 0,464 | L_7Am | 0,614 | L_7Am | 0,542 | | L_7Am | 0,289 | L_7Am | 0,308 | L_7Am | 0,223 |
| L_7PL | 0,492 | L_7PL | 0,605 | L_7PL | 0,494 | | L_7PL | 0,326 | L_7PL | 0,374 | L_7PL | 0,290 |
| L_7PC | 0,498 | L_7PC | 0,637 | L_7PC | 0,500 | | L_7PC | 0,345 | L_7PC | 0,429 | L_7PC | 0,185 |
| L_LIPv | 0,519 | L_LIPv | 0,660 | L_LIPv | 0,493 | | L_LIPv | 0,305 | L_LIPv | 0,347 | L_LIPv | 0,216 |
| L_VIP | 0,477 | L_VIP | 0,627 | L_VIP | 0,509 | | L_VIP | 0,343 | L_VIP | 0,312 | L_VIP | 0,182 |
| L_MIP | 0,554 | L_MIP | 0,722 | L_MIP | 0,526 | | L_MIP | 0,345 | L_MIP | 0,364 | L_MIP | 0,276 |
| L_1 | 0,528 | L_1 | 0,647 | L_1 | 0,480 | | L_1 | 0,387 | L_1 | 0,461 | L_1 | 0,360 |
| L_2 | 0,521 | L_2 | 0,635 | L_2 | 0,506 | | L_2 | 0,302 | L_2 | 0,348 | L_2 | 0,359 |
| L_3a | 0,605 | L_3a | 0,686 | L_3a | 0,517 | | L_3a | 0,331 | L_3a | 0,357 | L_3a | 0,376 |
| L_6d | 0,487 | L_6d | 0,661 | L_6d | 0,478 | | L_6d | 0,416 | L_6d | 0,413 | L_6d | 0,281 |

| | | | | | | | | | | | | |
|---|---|---|---|---|---|---|---|---|---|---|---|---|
| L_6mp | 0,523 | L_6mp | 0,687 | L_6mp | 0,457 | | L_6mp | 0,352 | L_6mp | 0,413 | L_6mp | 0,300 |
| L_6v | 0,449 | L_6v | 0,632 | L_6v | 0,544 | | L_6v | 0,271 | L_6v | 0,296 | L_6v | 0,259 |
| L_p24pr | 0,452 | L_p24pr | 0,532 | L_p24pr | 0,502 | | L_p24pr | 0,255 | L_p24pr | 0,181 | L_p24pr | 0,120 |
| L_33pr | 0,372 | L_33pr | 0,354 | L_33pr | 0,485 | | L_33pr | 0,245 | L_33pr | 0,159 | L_33pr | 0,179 |
| L_a24pr | 0,371 | L_a24pr | 0,497 | L_a24pr | 0,514 | | L_a24pr | 0,209 | L_a24pr | 0,219 | L_a24pr | 0,284 |
| L_p32pr | 0,356 | L_p32pr | 0,624 | L_p32pr | 0,598 | | L_p32pr | 0,204 | L_p32pr | 0,260 | L_p32pr | 0,296 |
| L_a24 | 0,350 | L_a24 | 0,516 | L_a24 | 0,507 | | L_a24 | 0,161 | L_a24 | 0,199 | L_a24 | 0,185 |
| L_d32 | 0,444 | L_d32 | 0,685 | L_d32 | 0,567 | | L_d32 | 0,331 | L_d32 | 0,367 | L_d32 | 0,281 |
| L_8BM | 0,480 | L_8BM | 0,700 | L_8BM | 0,569 | | L_8BM | 0,361 | L_8BM | 0,382 | L_8BM | 0,158 |
| L_p32 | 0,400 | L_p32 | 0,573 | L_p32 | 0,469 | | L_p32 | 0,217 | L_p32 | 0,289 | L_p32 | 0,202 |
| L_10r | 0,408 | L_10r | 0,481 | L_10r | 0,436 | | L_10r | 0,136 | L_10r | 0,213 | L_10r | 0,224 |
| L_47m | 0,494 | L_47m | 0,689 | L_47m | 0,488 | | L_47m | 0,300 | L_47m | 0,332 | L_47m | 0,120 |
| L_8Av | 0,453 | L_8Av | 0,653 | L_8Av | 0,495 | | L_8Av | 0,378 | L_8Av | 0,475 | L_8Av | 0,209 |
| L_8Ad | 0,491 | L_8Ad | 0,668 | L_8Ad | 0,470 | | L_8Ad | 0,282 | L_8Ad | 0,371 | L_8Ad | 0,181 |
| L_9m | 0,479 | L_9m | 0,671 | L_9m | 0,485 | | L_9m | 0,306 | L_9m | 0,342 | L_9m | 0,257 |
| L_8BL | 0,508 | L_8BL | 0,709 | L_8BL | 0,443 | | L_8BL | 0,377 | L_8BL | 0,313 | L_8BL | 0,234 |
| L_9p | 0,494 | L_9p | 0,687 | L_9p | 0,498 | | L_9p | 0,305 | L_9p | 0,336 | L_9p | 0,253 |
| L_10d | 0,428 | L_10d | 0,642 | L_10d | 0,373 | | L_10d | 0,302 | L_10d | 0,336 | L_10d | 0,278 |
| L_8C | 0,523 | L_8C | 0,699 | L_8C | 0,559 | | L_8C | 0,350 | L_8C | 0,369 | L_8C | 0,249 |
| L_44 | 0,492 | L_44 | 0,684 | L_44 | 0,598 | | L_44 | 0,366 | L_44 | 0,390 | L_44 | 0,190 |
| L_45 | 0,521 | L_45 | 0,680 | L_45 | 0,542 | | L_45 | 0,329 | L_45 | 0,349 | L_45 | 0,282 |
| L_47l | 0,560 | L_47l | 0,704 | L_47l | 0,508 | | L_47l | 0,279 | L_47l | 0,336 | L_47l | 0,195 |
| L_a47r | 0,479 | L_a47r | 0,639 | L_a47r | 0,433 | | L_a47r | 0,240 | L_a47r | 0,335 | L_a47r | 0,160 |
| L_6r | 0,533 | L_6r | 0,683 | L_6r | 0,629 | | L_6r | 0,244 | L_6r | 0,279 | L_6r | 0,190 |
| L_IFJa | 0,531 | L_IFJa | 0,708 | L_IFJa | 0,524 | | L_IFJa | 0,308 | L_IFJa | 0,363 | L_IFJa | 0,219 |
| L_IFJp | 0,507 | L_IFJp | 0,695 | L_IFJp | 0,582 | | L_IFJp | 0,354 | L_IFJp | 0,354 | L_IFJp | 0,261 |
| L_IFSp | 0,525 | L_IFSp | 0,712 | L_IFSp | 0,489 | | L_IFSp | 0,366 | L_IFSp | 0,347 | L_IFSp | 0,202 |
| L_IFSa | 0,464 | L_IFSa | 0,686 | L_IFSa | 0,527 | | L_IFSa | 0,276 | L_IFSa | 0,390 | L_IFSa | 0,161 |
| L_p9.46v | 0,472 | L_p9.46v | 0,650 | L_p9.46v | 0,521 | | L_p9.46v | 0,267 | L_p9.46v | 0,337 | L_p9.46v | 0,219 |
| L_46 | 0,425 | L_46 | 0,637 | L_46 | 0,487 | | L_46 | 0,189 | L_46 | 0,399 | L_46 | 0,228 |

| | | | | | | | | | | | | |
|---|---|---|---|---|---|---|---|---|---|---|---|---|
| L_a9.46v | 0,540 | L_a9.46v | 0,647 | L_a9.46v | 0,497 | | L_a9.46v | 0,203 | L_a9.46v | 0,339 | L_a9.46v | 0,261 |
| L_9.46d | 0,486 | L_9.46d | 0,649 | L_9.46d | 0,514 | | L_9.46d | 0,269 | L_9.46d | 0,377 | L_9.46d | 0,268 |
| L_9a | 0,502 | L_9a | 0,680 | L_9a | 0,503 | | L_9a | 0,363 | L_9a | 0,432 | L_9a | 0,308 |
| L_10v | 0,458 | L_10v | 0,419 | L_10v | 0,436 | | L_10v | 0,184 | L_10v | 0,103 | L_10v | 0,189 |
| L_a10p | 0,405 | L_a10p | 0,564 | L_a10p | 0,437 | | L_a10p | 0,221 | L_a10p | 0,270 | L_a10p | 0,236 |
| L_10pp | 0,502 | L_10pp | 0,378 | L_10pp | 0,391 | | L_10pp | 0,307 | L_10pp | 0,108 | L_10pp | 0,172 |
| L_11l | 0,450 | L_11l | 0,439 | L_11l | 0,388 | | L_11l | 0,198 | L_11l | 0,151 | L_11l | 0,227 |
| L_13l | 0,559 | L_13l | 0,507 | L_13l | 0,407 | | L_13l | 0,297 | L_13l | 0,024 | L_13l | 0,164 |
| L_OFC | 0,560 | L_OFC | 0,341 | L_OFC | 0,347 | | L_OFC | 0,299 | L_OFC | 0,073 | L_OFC | 0,135 |
| L_47s | 0,545 | L_47s | 0,677 | L_47s | 0,483 | | L_47s | 0,270 | L_47s | 0,299 | L_47s | 0,119 |
| L_LIPd | 0,564 | L_LIPd | 0,741 | L_LIPd | 0,501 | | L_LIPd | 0,342 | L_LIPd | 0,393 | L_LIPd | 0,197 |
| L_6a | 0,439 | L_6a | 0,653 | L_6a | 0,539 | | L_6a | 0,194 | L_6a | 0,271 | L_6a | 0,221 |
| L_i6.8 | 0,484 | L_i6.8 | 0,662 | L_i6.8 | 0,457 | | L_i6.8 | 0,302 | L_i6.8 | 0,351 | L_i6.8 | 0,111 |
| L_s6.8 | 0,466 | L_s6.8 | 0,676 | L_s6.8 | 0,513 | | L_s6.8 | 0,302 | L_s6.8 | 0,414 | L_s6.8 | 0,181 |
| L_43 | 0,569 | L_43 | 0,688 | L_43 | 0,544 | | L_43 | 0,233 | L_43 | 0,212 | L_43 | 0,282 |
| L_OP4 | 0,552 | L_OP4 | 0,668 | L_OP4 | 0,537 | | L_OP4 | 0,307 | L_OP4 | 0,319 | L_OP4 | 0,281 |
| L_OP1 | 0,589 | L_OP1 | 0,711 | L_OP1 | 0,572 | | L_OP1 | 0,319 | L_OP1 | 0,355 | L_OP1 | 0,307 |
| L_OP2.3 | 0,543 | L_OP2.3 | 0,662 | L_OP2.3 | 0,597 | | L_OP2.3 | 0,295 | L_OP2.3 | 0,411 | L_OP2.3 | 0,257 |
| L_52 | 0,448 | L_52 | 0,600 | L_52 | 0,457 | | L_52 | 0,178 | L_52 | 0,241 | L_52 | 0,200 |
| L_RI | 0,518 | L_RI | 0,696 | L_RI | 0,577 | | L_RI | 0,228 | L_RI | 0,323 | L_RI | 0,320 |
| L_PFcm | 0,589 | L_PFcm | 0,695 | L_PFcm | 0,567 | | L_PFcm | 0,276 | L_PFcm | 0,270 | L_PFcm | 0,229 |
| L_PoI2 | 0,472 | L_PoI2 | 0,580 | L_PoI2 | 0,495 | | L_PoI2 | 0,213 | L_PoI2 | 0,164 | L_PoI2 | 0,176 |
| L_TA2 | 0,500 | L_TA2 | 0,568 | L_TA2 | 0,415 | | L_TA2 | 0,237 | L_TA2 | 0,250 | L_TA2 | 0,160 |
| L_FOP4 | 0,469 | L_FOP4 | 0,684 | L_FOP4 | 0,606 | | L_FOP4 | 0,241 | L_FOP4 | 0,254 | L_FOP4 | 0,287 |
| L_MI | 0,465 | L_MI | 0,567 | L_MI | 0,553 | | L_MI | 0,237 | L_MI | 0,205 | L_MI | 0,241 |
| L_Pir | 0,496 | L_Pir | 0,448 | L_Pir | 0,353 | | L_Pir | 0,380 | L_Pir | 0,165 | L_Pir | 0,031 |
| L_AVI | 0,488 | L_AVI | 0,617 | L_AVI | 0,552 | | L_AVI | 0,394 | L_AVI | 0,184 | L_AVI | 0,242 |
| L_AAIC | 0,556 | L_AAIC | 0,543 | L_AAIC | 0,525 | | L_AAIC | 0,371 | L_AAIC | 0,127 | L_AAIC | 0,081 |
| L_FOP1 | 0,557 | L_FOP1 | 0,682 | L_FOP1 | 0,623 | | L_FOP1 | 0,281 | L_FOP1 | 0,320 | L_FOP1 | 0,261 |
| L_FOP3 | 0,372 | L_FOP3 | 0,529 | L_FOP3 | 0,620 | | L_FOP3 | 0,193 | L_FOP3 | 0,278 | L_FOP3 | 0,317 |

| | | | | | | | | | | | | |
|---|---|---|---|---|---|---|---|---|---|---|---|---|
| L_FOP2 | 0,409 | L_FOP2 | 0,594 | L_FOP2 | 0,609 | | L_FOP2 | 0,209 | L_FOP2 | 0,326 | L_FOP2 | 0,304 |
| L_PFt | 0,446 | L_PFt | 0,641 | L_PFt | 0,517 | | L_PFt | 0,351 | L_PFt | 0,374 | L_PFt | 0,388 |
| L_AIP | 0,547 | L_AIP | 0,679 | L_AIP | 0,498 | | L_AIP | 0,290 | L_AIP | 0,290 | L_AIP | 0,277 |
| L_EC | 0,449 | L_EC | 0,379 | L_EC | 0,417 | | L_EC | 0,307 | L_EC | 0,087 | L_EC | 0,223 |
| L_PreS | 0,452 | L_PreS | 0,519 | L_PreS | 0,448 | | L_PreS | 0,222 | L_PreS | 0,185 | L_PreS | 0,176 |
| L_H | 0,390 | L_H | 0,342 | L_H | 0,353 | | L_H | 0,221 | L_H | 0,127 | L_H | 0,205 |
| L_ProS | 0,543 | L_ProS | 0,685 | L_ProS | 0,424 | | L_ProS | 0,383 | L_ProS | 0,436 | L_ProS | 0,268 |
| L_PeEc | 0,524 | L_PeEc | 0,406 | L_PeEc | 0,467 | | L_PeEc | 0,279 | L_PeEc | 0,067 | L_PeEc | 0,115 |
| L_STGa | 0,543 | L_STGa | 0,600 | L_STGa | 0,464 | | L_STGa | 0,181 | L_STGa | 0,250 | L_STGa | 0,210 |
| L_PBelt | 0,568 | L_PBelt | 0,694 | L_PBelt | 0,532 | | L_PBelt | 0,243 | L_PBelt | 0,365 | L_PBelt | 0,296 |
| L_A5 | 0,532 | L_A5 | 0,687 | L_A5 | 0,538 | | L_A5 | 0,234 | L_A5 | 0,245 | L_A5 | 0,293 |
| L_PHA1 | 0,486 | L_PHA1 | 0,497 | L_PHA1 | 0,448 | | L_PHA1 | 0,222 | L_PHA1 | 0,201 | L_PHA1 | 0,121 |
| L_PHA3 | 0,434 | L_PHA3 | 0,476 | L_PHA3 | 0,494 | | L_PHA3 | 0,291 | L_PHA3 | 0,132 | L_PHA3 | 0,152 |
| L_STSda | 0,520 | L_STSda | 0,655 | L_STSda | 0,550 | | L_STSda | 0,237 | L_STSda | 0,251 | L_STSda | 0,318 |
| L_STSdp | 0,493 | L_STSdp | 0,654 | L_STSdp | 0,503 | | L_STSdp | 0,326 | L_STSdp | 0,332 | L_STSdp | 0,309 |
| L_STSvp | 0,496 | L_STSvp | 0,683 | L_STSvp | 0,487 | | L_STSvp | 0,214 | L_STSvp | 0,354 | L_STSvp | 0,273 |
| L_TGd | 0,507 | L_TGd | 0,540 | L_TGd | 0,408 | | L_TGd | 0,322 | L_TGd | 0,099 | L_TGd | 0,073 |
| L_TE1a | 0,543 | L_TE1a | 0,626 | L_TE1a | 0,444 | | L_TE1a | 0,144 | L_TE1a | 0,237 | L_TE1a | 0,126 |
| L_TE1p | 0,447 | L_TE1p | 0,625 | L_TE1p | 0,461 | | L_TE1p | 0,257 | L_TE1p | 0,328 | L_TE1p | 0,195 |
| L_TE2a | 0,540 | L_TE2a | 0,629 | L_TE2a | 0,412 | | L_TE2a | 0,317 | L_TE2a | 0,306 | L_TE2a | 0,083 |
| L_TF | 0,539 | L_TF | 0,488 | L_TF | 0,475 | | L_TF | 0,316 | L_TF | 0,154 | L_TF | 0,139 |
| L_TE2p | 0,512 | L_TE2p | 0,550 | L_TE2p | 0,467 | | L_TE2p | 0,175 | L_TE2p | 0,243 | L_TE2p | 0,114 |
| L_PHT | 0,532 | L_PHT | 0,600 | L_PHT | 0,487 | | L_PHT | 0,252 | L_PHT | 0,364 | L_PHT | 0,195 |
| L_PH | 0,437 | L_PH | 0,616 | L_PH | 0,488 | | L_PH | 0,263 | L_PH | 0,266 | L_PH | 0,227 |
| L_TPOJ1 | 0,552 | L_TPOJ1 | 0,710 | L_TPOJ1 | 0,535 | | L_TPOJ1 | 0,205 | L_TPOJ1 | 0,307 | L_TPOJ1 | 0,290 |
| L_TPOJ2 | 0,468 | L_TPOJ2 | 0,678 | L_TPOJ2 | 0,495 | | L_TPOJ2 | 0,283 | L_TPOJ2 | 0,342 | L_TPOJ2 | 0,241 |
| L_TPOJ3 | 0,547 | L_TPOJ3 | 0,755 | L_TPOJ3 | 0,460 | | L_TPOJ3 | 0,326 | L_TPOJ3 | 0,377 | L_TPOJ3 | 0,237 |
| L_DVT | 0,492 | L_DVT | 0,633 | L_DVT | 0,446 | | L_DVT | 0,283 | L_DVT | 0,321 | L_DVT | 0,216 |
| L_PGp | 0,422 | L_PGp | 0,592 | L_PGp | 0,413 | | L_PGp | 0,282 | L_PGp | 0,383 | L_PGp | 0,243 |
| L_IP2 | 0,542 | L_IP2 | 0,695 | L_IP2 | 0,492 | | L_IP2 | 0,276 | L_IP2 | 0,348 | L_IP2 | 0,199 |

| | | | | | | | | | | | | |
|---|---|---|---|---|---|---|---|---|---|---|---|---|
| L_IP1 | 0,479 | L_IP1 | 0,661 | L_IP1 | 0,482 | | L_IP1 | 0,345 | L_IP1 | 0,315 | L_IP1 | 0,233 |
| L_IP0 | 0,408 | L_IP0 | 0,644 | L_IP0 | 0,447 | | L_IP0 | 0,324 | L_IP0 | 0,304 | L_IP0 | 0,301 |
| L_PFop | 0,533 | L_PFop | 0,676 | L_PFop | 0,555 | | L_PFop | 0,320 | L_PFop | 0,378 | L_PFop | 0,215 |
| L_PF | 0,483 | L_PF | 0,643 | L_PF | 0,530 | | L_PF | 0,343 | L_PF | 0,376 | L_PF | 0,275 |
| L_PFm | 0,497 | L_PFm | 0,669 | L_PFm | 0,505 | | L_PFm | 0,376 | L_PFm | 0,447 | L_PFm | 0,232 |
| L_PGi | 0,521 | L_PGi | 0,680 | L_PGi | 0,470 | | L_PGi | 0,427 | L_PGi | 0,491 | L_PGi | 0,235 |
| L_PGs | 0,547 | L_PGs | 0,630 | L_PGs | 0,444 | | L_PGs | 0,385 | L_PGs | 0,426 | L_PGs | 0,261 |
| L_V6A | 0,468 | L_V6A | 0,597 | L_V6A | 0,476 | | L_V6A | 0,407 | L_V6A | 0,361 | L_V6A | 0,247 |
| L_VMV1 | 0,486 | L_VMV1 | 0,652 | L_VMV1 | 0,468 | | L_VMV1 | 0,320 | L_VMV1 | 0,444 | L_VMV1 | 0,251 |
| L_VMV3 | 0,431 | L_VMV3 | 0,579 | L_VMV3 | 0,461 | | L_VMV3 | 0,267 | L_VMV3 | 0,386 | L_VMV3 | 0,275 |
| L_PHA2 | 0,457 | L_PHA2 | 0,503 | L_PHA2 | 0,501 | | L_PHA2 | 0,376 | L_PHA2 | 0,250 | L_PHA2 | 0,214 |
| L_V4t | 0,506 | L_V4t | 0,672 | L_V4t | 0,435 | | L_V4t | 0,293 | L_V4t | 0,359 | L_V4t | 0,229 |
| L_FST | 0,490 | L_FST | 0,651 | L_FST | 0,469 | | L_FST | 0,287 | L_FST | 0,354 | L_FST | 0,237 |
| L_V3CD | 0,477 | L_V3CD | 0,628 | L_V3CD | 0,477 | | L_V3CD | 0,415 | L_V3CD | 0,450 | L_V3CD | 0,417 |
| L_LO3 | 0,426 | L_LO3 | 0,640 | L_LO3 | 0,466 | | L_LO3 | 0,279 | L_LO3 | 0,395 | L_LO3 | 0,265 |
| L_VMV2 | 0,440 | L_VMV2 | 0,596 | L_VMV2 | 0,481 | | L_VMV2 | 0,270 | L_VMV2 | 0,348 | L_VMV2 | 0,244 |
| L_31pd | 0,440 | L_31pd | 0,700 | L_31pd | 0,523 | | L_31pd | 0,367 | L_31pd | 0,414 | L_31pd | 0,278 |
| L_31a | 0,377 | L_31a | 0,629 | L_31a | 0,521 | | L_31a | 0,197 | L_31a | 0,309 | L_31a | 0,348 |
| L_VVC | 0,444 | L_VVC | 0,584 | L_VVC | 0,469 | | L_VVC | 0,268 | L_VVC | 0,311 | L_VVC | 0,273 |
| L_25 | 0,474 | L_25 | 0,363 | L_25 | 0,445 | | L_25 | 0,339 | L_25 | 0,141 | L_25 | 0,167 |
| L_s32 | 0,495 | L_s32 | 0,440 | L_s32 | 0,468 | | L_s32 | 0,376 | L_s32 | 0,175 | L_s32 | 0,261 |
| L_pOFC | 0,520 | L_pOFC | 0,359 | L_pOFC | 0,456 | | L_pOFC | 0,381 | L_pOFC | 0,069 | L_pOFC | 0,120 |
| L_PoI1 | 0,434 | L_PoI1 | 0,477 | L_PoI1 | 0,448 | | L_PoI1 | 0,185 | L_PoI1 | 0,124 | L_PoI1 | 0,140 |
| L_Ig | 0,479 | L_Ig | 0,648 | L_Ig | 0,530 | | L_Ig | 0,232 | L_Ig | 0,359 | L_Ig | 0,289 |
| L_FOP5 | 0,507 | L_FOP5 | 0,663 | L_FOP5 | 0,568 | | L_FOP5 | 0,376 | L_FOP5 | 0,340 | L_FOP5 | 0,237 |
| L_p10p | 0,394 | L_p10p | 0,610 | L_p10p | 0,395 | | L_p10p | 0,159 | L_p10p | 0,272 | L_p10p | 0,253 |
| L_p47r | 0,531 | L_p47r | 0,690 | L_p47r | 0,502 | | L_p47r | 0,276 | L_p47r | 0,285 | L_p47r | 0,229 |
| L_TGv | 0,526 | L_TGv | 0,513 | L_TGv | 0,503 | | L_TGv | 0,261 | L_TGv | 0,114 | L_TGv | 0,185 |
| L_MBelt | 0,533 | L_MBelt | 0,632 | L_MBelt | 0,498 | | L_MBelt | 0,227 | L_MBelt | 0,313 | L_MBelt | 0,227 |
| L_LBelt | 0,590 | L_LBelt | 0,720 | L_LBelt | 0,563 | | L_LBelt | 0,271 | L_LBelt | 0,386 | L_LBelt | 0,350 |

| | | | | | | | | | | | | |
|---|---|---|---|---|---|---|---|---|---|---|---|---|
| L_A4 | 0,553 | L_A4 | 0,669 | L_A4 | 0,514 | | L_A4 | 0,198 | L_A4 | 0,322 | L_A4 | 0,306 |
| L_STSva | 0,542 | L_STSva | 0,661 | L_STSva | 0,483 | | L_STSva | 0,213 | L_STSva | 0,330 | L_STSva | 0,186 |
| L_TE1m | 0,485 | L_TE1m | 0,664 | L_TE1m | 0,451 | | L_TE1m | 0,329 | L_TE1m | 0,228 | L_TE1m | 0,205 |
| L_PI | 0,463 | L_PI | 0,464 | L_PI | 0,136 | | L_PI | 0,361 | L_PI | 0,180 | L_PI | 0,053 |
| L_a32pr | 0,402 | L_a32pr | 0,665 | L_a32pr | 0,599 | | L_a32pr | 0,260 | L_a32pr | 0,250 | L_a32pr | 0,226 |
| L_p24 | 0,370 | L_p24 | 0,507 | L_p24 | 0,521 | | L_p24 | 0,202 | L_p24 | 0,126 | L_p24 | 0,283 |
| R_V1 | 0,435 | R_V1 | 0,583 | R_V1 | 0,420 | | R_V1 | 0,339 | R_V1 | 0,386 | R_V1 | 0,349 |
| R_MST | 0,521 | R_MST | 0,652 | R_MST | 0,473 | | R_MST | 0,310 | R_MST | 0,400 | R_MST | 0,309 |
| R_V6 | 0,494 | R_V6 | 0,672 | R_V6 | 0,424 | | R_V6 | 0,414 | R_V6 | 0,381 | R_V6 | 0,240 |
| R_V2 | 0,449 | R_V2 | 0,637 | R_V2 | 0,435 | | R_V2 | 0,392 | R_V2 | 0,349 | R_V2 | 0,373 |
| R_V3 | 0,468 | R_V3 | 0,595 | R_V3 | 0,403 | | R_V3 | 0,511 | R_V3 | 0,457 | R_V3 | 0,377 |
| R_V4 | 0,449 | R_V4 | 0,598 | R_V4 | 0,434 | | R_V4 | 0,462 | R_V4 | 0,506 | R_V4 | 0,342 |
| R_V8 | 0,469 | R_V8 | 0,613 | R_V8 | 0,436 | | R_V8 | 0,275 | R_V8 | 0,301 | R_V8 | 0,278 |
| R_4 | 0,549 | R_4 | 0,692 | R_4 | 0,509 | | R_4 | 0,356 | R_4 | 0,387 | R_4 | 0,349 |
| R_3b | 0,556 | R_3b | 0,659 | R_3b | 0,491 | | R_3b | 0,286 | R_3b | 0,381 | R_3b | 0,314 |
| R_FEF | 0,524 | R_FEF | 0,641 | R_FEF | 0,530 | | R_FEF | 0,249 | R_FEF | 0,259 | R_FEF | 0,216 |
| R_PEF | 0,508 | R_PEF | 0,711 | R_PEF | 0,540 | | R_PEF | 0,277 | R_PEF | 0,342 | R_PEF | 0,225 |
| R_55b | 0,549 | R_55b | 0,672 | R_55b | 0,516 | | R_55b | 0,244 | R_55b | 0,289 | R_55b | 0,099 |
| R_V3A | 0,464 | R_V3A | 0,641 | R_V3A | 0,429 | | R_V3A | 0,472 | R_V3A | 0,498 | R_V3A | 0,381 |
| R_RSC | 0,411 | R_RSC | 0,527 | R_RSC | 0,532 | | R_RSC | 0,211 | R_RSC | 0,199 | R_RSC | 0,312 |
| R_POS2 | 0,468 | R_POS2 | 0,623 | R_POS2 | 0,547 | | R_POS2 | 0,311 | R_POS2 | 0,299 | R_POS2 | 0,272 |
| R_V7 | 0,501 | R_V7 | 0,649 | R_V7 | 0,537 | | R_V7 | 0,416 | R_V7 | 0,408 | R_V7 | 0,241 |
| R_IPS1 | 0,488 | R_IPS1 | 0,680 | R_IPS1 | 0,483 | | R_IPS1 | 0,358 | R_IPS1 | 0,315 | R_IPS1 | 0,249 |
| R_FFC | 0,397 | R_FFC | 0,557 | R_FFC | 0,464 | | R_FFC | 0,177 | R_FFC | 0,178 | R_FFC | 0,262 |
| R_V3B | 0,479 | R_V3B | 0,679 | R_V3B | 0,495 | | R_V3B | 0,451 | R_V3B | 0,455 | R_V3B | 0,301 |
| R_LO1 | 0,458 | R_LO1 | 0,646 | R_LO1 | 0,417 | | R_LO1 | 0,398 | R_LO1 | 0,435 | R_LO1 | 0,274 |
| R_LO2 | 0,410 | R_LO2 | 0,545 | R_LO2 | 0,421 | | R_LO2 | 0,374 | R_LO2 | 0,367 | R_LO2 | 0,327 |
| R_PIT | 0,376 | R_PIT | 0,528 | R_PIT | 0,411 | | R_PIT | 0,276 | R_PIT | 0,307 | R_PIT | 0,239 |
| R_MT | 0,520 | R_MT | 0,670 | R_MT | 0,453 | | R_MT | 0,268 | R_MT | 0,402 | R_MT | 0,331 |
| R_A1 | 0,524 | R_A1 | 0,667 | R_A1 | 0,438 | | R_A1 | 0,352 | R_A1 | 0,389 | R_A1 | 0,145 |

| | | | | | | | | | | | | |
|---|---|---|---|---|---|---|---|---|---|---|---|---|
| R_PSL | 0,438 | R_PSL | 0,622 | R_PSL | 0,496 | | R_PSL | 0,358 | R_PSL | 0,402 | R_PSL | 0,263 |
| R_SFL | 0,417 | R_SFL | 0,655 | R_SFL | 0,463 | | R_SFL | 0,267 | R_SFL | 0,288 | R_SFL | 0,159 |
| R_PCV | 0,442 | R_PCV | 0,638 | R_PCV | 0,550 | | R_PCV | 0,229 | R_PCV | 0,297 | R_PCV | 0,249 |
| R_STV | 0,501 | R_STV | 0,687 | R_STV | 0,526 | | R_STV | 0,235 | R_STV | 0,261 | R_STV | 0,200 |
| R_7Pm | 0,488 | R_7Pm | 0,636 | R_7Pm | 0,529 | | R_7Pm | 0,310 | R_7Pm | 0,273 | R_7Pm | 0,166 |
| R_7m | 0,404 | R_7m | 0,650 | R_7m | 0,478 | | R_7m | 0,342 | R_7m | 0,300 | R_7m | 0,394 |
| R_POS1 | 0,403 | R_POS1 | 0,547 | R_POS1 | 0,422 | | R_POS1 | 0,252 | R_POS1 | 0,197 | R_POS1 | 0,189 |
| R_23d | 0,325 | R_23d | 0,495 | R_23d | 0,511 | | R_23d | 0,178 | R_23d | 0,173 | R_23d | 0,247 |
| R_v23ab | 0,452 | R_v23ab | 0,595 | R_v23ab | 0,533 | | R_v23ab | 0,287 | R_v23ab | 0,251 | R_v23ab | 0,266 |
| R_d23ab | 0,440 | R_d23ab | 0,550 | R_d23ab | 0,522 | | R_d23ab | 0,225 | R_d23ab | 0,307 | R_d23ab | 0,317 |
| R_31pv | 0,495 | R_31pv | 0,696 | R_31pv | 0,504 | | R_31pv | 0,312 | R_31pv | 0,321 | R_31pv | 0,251 |
| R_5m | 0,541 | R_5m | 0,661 | R_5m | 0,516 | | R_5m | 0,397 | R_5m | 0,397 | R_5m | 0,362 |
| R_5mv | 0,512 | R_5mv | 0,672 | R_5mv | 0,585 | | R_5mv | 0,273 | R_5mv | 0,330 | R_5mv | 0,156 |
| R_23c | 0,418 | R_23c | 0,651 | R_23c | 0,525 | | R_23c | 0,224 | R_23c | 0,220 | R_23c | 0,159 |
| R_5L | 0,488 | R_5L | 0,656 | R_5L | 0,490 | | R_5L | 0,312 | R_5L | 0,355 | R_5L | 0,239 |
| R_24dd | 0,512 | R_24dd | 0,684 | R_24dd | 0,589 | | R_24dd | 0,275 | R_24dd | 0,329 | R_24dd | 0,300 |
| R_24dv | 0,499 | R_24dv | 0,647 | R_24dv | 0,597 | | R_24dv | 0,208 | R_24dv | 0,246 | R_24dv | 0,288 |
| R_7AL | 0,478 | R_7AL | 0,663 | R_7AL | 0,461 | | R_7AL | 0,304 | R_7AL | 0,270 | R_7AL | 0,218 |
| R_SCEF | 0,442 | R_SCEF | 0,680 | R_SCEF | 0,541 | | R_SCEF | 0,205 | R_SCEF | 0,212 | R_SCEF | 0,257 |
| R_6ma | 0,452 | R_6ma | 0,629 | R_6ma | 0,537 | | R_6ma | 0,269 | R_6ma | 0,372 | R_6ma | 0,185 |
| R_7Am | 0,498 | R_7Am | 0,612 | R_7Am | 0,490 | | R_7Am | 0,350 | R_7Am | 0,330 | R_7Am | 0,187 |
| R_7PL | 0,508 | R_7PL | 0,721 | R_7PL | 0,538 | | R_7PL | 0,321 | R_7PL | 0,352 | R_7PL | 0,249 |
| R_7PC | 0,500 | R_7PC | 0,624 | R_7PC | 0,511 | | R_7PC | 0,348 | R_7PC | 0,386 | R_7PC | 0,257 |
| R_LIPv | 0,500 | R_LIPv | 0,705 | R_LIPv | 0,465 | | R_LIPv | 0,275 | R_LIPv | 0,310 | R_LIPv | 0,233 |
| R_VIP | 0,519 | R_VIP | 0,630 | R_VIP | 0,467 | | R_VIP | 0,368 | R_VIP | 0,370 | R_VIP | 0,207 |
| R_MIP | 0,498 | R_MIP | 0,688 | R_MIP | 0,527 | | R_MIP | 0,352 | R_MIP | 0,325 | R_MIP | 0,248 |
| R_1 | 0,578 | R_1 | 0,642 | R_1 | 0,463 | | R_1 | 0,410 | R_1 | 0,484 | R_1 | 0,372 |
| R_2 | 0,521 | R_2 | 0,653 | R_2 | 0,474 | | R_2 | 0,340 | R_2 | 0,400 | R_2 | 0,348 |
| R_3a | 0,566 | R_3a | 0,690 | R_3a | 0,518 | | R_3a | 0,356 | R_3a | 0,285 | R_3a | 0,344 |
| R_6d | 0,559 | R_6d | 0,647 | R_6d | 0,508 | | R_6d | 0,382 | R_6d | 0,418 | R_6d | 0,296 |

| | | | | | | | | | | | | |
|---|---|---|---|---|---|---|---|---|---|---|---|---|
| R_6mp | 0,494 | R_6mp | 0,688 | R_6mp | 0,496 | | R_6mp | 0,291 | R_6mp | 0,336 | R_6mp | 0,215 |
| R_6v | 0,544 | R_6v | 0,666 | R_6v | 0,485 | | R_6v | 0,303 | R_6v | 0,292 | R_6v | 0,288 |
| R_p24pr | 0,385 | R_p24pr | 0,564 | R_p24pr | 0,517 | | R_p24pr | 0,135 | R_p24pr | 0,290 | R_p24pr | 0,187 |
| R_33pr | 0,450 | R_33pr | 0,391 | R_33pr | 0,528 | | R_33pr | 0,276 | R_33pr | 0,166 | R_33pr | 0,215 |
| R_a24pr | 0,382 | R_a24pr | 0,476 | R_a24pr | 0,533 | | R_a24pr | 0,189 | R_a24pr | 0,249 | R_a24pr | 0,162 |
| R_p32pr | 0,411 | R_p32pr | 0,656 | R_p32pr | 0,588 | | R_p32pr | 0,161 | R_p32pr | 0,259 | R_p32pr | 0,282 |
| R_a24 | 0,448 | R_a24 | 0,511 | R_a24 | 0,500 | | R_a24 | 0,301 | R_a24 | 0,193 | R_a24 | 0,320 |
| R_d32 | 0,402 | R_d32 | 0,705 | R_d32 | 0,563 | | R_d32 | 0,236 | R_d32 | 0,339 | R_d32 | 0,265 |
| R_8BM | 0,414 | R_8BM | 0,662 | R_8BM | 0,594 | | R_8BM | 0,236 | R_8BM | 0,344 | R_8BM | 0,235 |
| R_p32 | 0,424 | R_p32 | 0,509 | R_p32 | 0,515 | | R_p32 | 0,309 | R_p32 | 0,246 | R_p32 | 0,280 |
| R_10r | 0,376 | R_10r | 0,456 | R_10r | 0,500 | | R_10r | 0,158 | R_10r | 0,145 | R_10r | 0,203 |
| R_47m | 0,521 | R_47m | 0,591 | R_47m | 0,433 | | R_47m | 0,258 | R_47m | 0,238 | R_47m | 0,152 |
| R_8Av | 0,530 | R_8Av | 0,648 | R_8Av | 0,468 | | R_8Av | 0,370 | R_8Av | 0,452 | R_8Av | 0,223 |
| R_8Ad | 0,437 | R_8Ad | 0,652 | R_8Ad | 0,486 | | R_8Ad | 0,276 | R_8Ad | 0,369 | R_8Ad | 0,265 |
| R_9m | 0,423 | R_9m | 0,640 | R_9m | 0,499 | | R_9m | 0,267 | R_9m | 0,276 | R_9m | 0,253 |
| R_8BL | 0,428 | R_8BL | 0,671 | R_8BL | 0,467 | | R_8BL | 0,319 | R_8BL | 0,362 | R_8BL | 0,210 |
| R_9p | 0,466 | R_9p | 0,642 | R_9p | 0,487 | | R_9p | 0,276 | R_9p | 0,384 | R_9p | 0,291 |
| R_10d | 0,387 | R_10d | 0,618 | R_10d | 0,420 | | R_10d | 0,192 | R_10d | 0,220 | R_10d | 0,327 |
| R_8C | 0,487 | R_8C | 0,666 | R_8C | 0,554 | | R_8C | 0,368 | R_8C | 0,449 | R_8C | 0,244 |
| R_44 | 0,480 | R_44 | 0,652 | R_44 | 0,594 | | R_44 | 0,343 | R_44 | 0,305 | R_44 | 0,235 |
| R_45 | 0,476 | R_45 | 0,678 | R_45 | 0,555 | | R_45 | 0,236 | R_45 | 0,209 | R_45 | 0,194 |
| R_47l | 0,512 | R_47l | 0,632 | R_47l | 0,472 | | R_47l | 0,230 | R_47l | 0,214 | R_47l | 0,184 |
| R_a47r | 0,435 | R_a47r | 0,613 | R_a47r | 0,407 | | R_a47r | 0,241 | R_a47r | 0,298 | R_a47r | 0,177 |
| R_6r | 0,525 | R_6r | 0,648 | R_6r | 0,554 | | R_6r | 0,307 | R_6r | 0,280 | R_6r | 0,293 |
| R_IFJa | 0,507 | R_IFJa | 0,726 | R_IFJa | 0,500 | | R_IFJa | 0,223 | R_IFJa | 0,389 | R_IFJa | 0,204 |
| R_IFJp | 0,506 | R_IFJp | 0,682 | R_IFJp | 0,478 | | R_IFJp | 0,250 | R_IFJp | 0,314 | R_IFJp | 0,199 |
| R_IFSp | 0,506 | R_IFSp | 0,676 | R_IFSp | 0,469 | | R_IFSp | 0,326 | R_IFSp | 0,369 | R_IFSp | 0,165 |
| R_IFSa | 0,516 | R_IFSa | 0,656 | R_IFSa | 0,468 | | R_IFSa | 0,232 | R_IFSa | 0,304 | R_IFSa | 0,153 |
| R_p9.46v | 0,490 | R_p9.46v | 0,661 | R_p9.46v | 0,511 | | R_p9.46v | 0,464 | R_p9.46v | 0,478 | R_p9.46v | 0,223 |
| R_46 | 0,452 | R_46 | 0,650 | R_46 | 0,505 | | R_46 | 0,324 | R_46 | 0,348 | R_46 | 0,228 |

| | | | | | | | | | | | | |
|---|---|---|---|---|---|---|---|---|---|---|---|---|
| R_a9.46v | 0,434 | R_a9.46v | 0,607 | R_a9.46v | 0,475 | | R_a9.46v | 0,276 | R_a9.46v | 0,380 | R_a9.46v | 0,276 |
| R_9.46d | 0,454 | R_9.46d | 0,621 | R_9.46d | 0,472 | | R_9.46d | 0,288 | R_9.46d | 0,327 | R_9.46d | 0,347 |
| R_9a | 0,458 | R_9a | 0,667 | R_9a | 0,491 | | R_9a | 0,248 | R_9a | 0,366 | R_9a | 0,285 |
| R_10v | 0,447 | R_10v | 0,448 | R_10v | 0,479 | | R_10v | 0,216 | R_10v | 0,074 | R_10v | 0,211 |
| R_a10p | 0,373 | R_a10p | 0,501 | R_a10p | 0,385 | | R_a10p | 0,193 | R_a10p | 0,245 | R_a10p | 0,266 |
| R_10pp | 0,531 | R_10pp | 0,500 | R_10pp | 0,434 | | R_10pp | 0,247 | R_10pp | 0,188 | R_10pp | 0,201 |
| R_11l | 0,401 | R_11l | 0,465 | R_11l | 0,363 | | R_11l | 0,156 | R_11l | 0,096 | R_11l | 0,193 |
| R_13l | 0,549 | R_13l | 0,337 | R_13l | 0,411 | | R_13l | 0,322 | R_13l | 0,054 | R_13l | 0,094 |
| R_OFC | 0,500 | R_OFC | 0,290 | R_OFC | 0,411 | | R_OFC | 0,319 | R_OFC | 0,043 | R_OFC | 0,185 |
| R_47s | 0,565 | R_47s | 0,607 | R_47s | 0,451 | | R_47s | 0,192 | R_47s | 0,190 | R_47s | 0,152 |
| R_LIPd | 0,511 | R_LIPd | 0,713 | R_LIPd | 0,469 | | R_LIPd | 0,306 | R_LIPd | 0,332 | R_LIPd | 0,197 |
| R_6a | 0,466 | R_6a | 0,665 | R_6a | 0,505 | | R_6a | 0,284 | R_6a | 0,298 | R_6a | 0,279 |
| R_i6.8 | 0,529 | R_i6.8 | 0,634 | R_i6.8 | 0,467 | | R_i6.8 | 0,340 | R_i6.8 | 0,411 | R_i6.8 | 0,201 |
| R_s6.8 | 0,448 | R_s6.8 | 0,685 | R_s6.8 | 0,476 | | R_s6.8 | 0,260 | R_s6.8 | 0,333 | R_s6.8 | 0,137 |
| R_43 | 0,528 | R_43 | 0,666 | R_43 | 0,568 | | R_43 | 0,254 | R_43 | 0,227 | R_43 | 0,189 |
| R_OP4 | 0,529 | R_OP4 | 0,681 | R_OP4 | 0,573 | | R_OP4 | 0,385 | R_OP4 | 0,408 | R_OP4 | 0,327 |
| R_OP1 | 0,576 | R_OP1 | 0,701 | R_OP1 | 0,546 | | R_OP1 | 0,314 | R_OP1 | 0,379 | R_OP1 | 0,343 |
| R_OP2.3 | 0,529 | R_OP2.3 | 0,695 | R_OP2.3 | 0,591 | | R_OP2.3 | 0,321 | R_OP2.3 | 0,420 | R_OP2.3 | 0,342 |
| R_52 | 0,451 | R_52 | 0,623 | R_52 | 0,418 | | R_52 | 0,227 | R_52 | 0,354 | R_52 | 0,171 |
| R_RI | 0,535 | R_RI | 0,694 | R_RI | 0,577 | | R_RI | 0,255 | R_RI | 0,394 | R_RI | 0,242 |
| R_PFcm | 0,549 | R_PFcm | 0,701 | R_PFcm | 0,551 | | R_PFcm | 0,261 | R_PFcm | 0,316 | R_PFcm | 0,290 |
| R_PoI2 | 0,463 | R_PoI2 | 0,647 | R_PoI2 | 0,506 | | R_PoI2 | 0,160 | R_PoI2 | 0,154 | R_PoI2 | 0,214 |
| R_TA2 | 0,434 | R_TA2 | 0,597 | R_TA2 | 0,440 | | R_TA2 | 0,205 | R_TA2 | 0,250 | R_TA2 | 0,211 |
| R_FOP4 | 0,468 | R_FOP4 | 0,635 | R_FOP4 | 0,613 | | R_FOP4 | 0,225 | R_FOP4 | 0,269 | R_FOP4 | 0,303 |
| R_MI | 0,447 | R_MI | 0,565 | R_MI | 0,533 | | R_MI | 0,244 | R_MI | 0,237 | R_MI | 0,305 |
| R_Pir | 0,477 | R_Pir | 0,507 | R_Pir | 0,396 | | R_Pir | 0,363 | R_Pir | 0,212 | R_Pir | 0,013 |
| R_AVI | 0,473 | R_AVI | 0,631 | R_AVI | 0,548 | | R_AVI | 0,342 | R_AVI | 0,240 | R_AVI | 0,270 |
| R_AAIC | 0,474 | R_AAIC | 0,592 | R_AAIC | 0,514 | | R_AAIC | 0,332 | R_AAIC | 0,154 | R_AAIC | 0,126 |
| R_FOP1 | 0,527 | R_FOP1 | 0,673 | R_FOP1 | 0,620 | | R_FOP1 | 0,254 | R_FOP1 | 0,311 | R_FOP1 | 0,277 |
| R_FOP3 | 0,462 | R_FOP3 | 0,690 | R_FOP3 | 0,600 | | R_FOP3 | 0,276 | R_FOP3 | 0,345 | R_FOP3 | 0,349 |

| | | | | | | | | | | | | |
|---|---|---|---|---|---|---|---|---|---|---|---|---|
| R_FOP2 | 0,505 | R_FOP2 | 0,687 | R_FOP2 | 0,592 | | R_FOP2 | 0,265 | R_FOP2 | 0,454 | R_FOP2 | 0,351 |
| R_PFt | 0,546 | R_PFt | 0,659 | R_PFt | 0,497 | | R_PFt | 0,384 | R_PFt | 0,324 | R_PFt | 0,289 |
| R_AIP | 0,533 | R_AIP | 0,654 | R_AIP | 0,491 | | R_AIP | 0,415 | R_AIP | 0,375 | R_AIP | 0,279 |
| R_EC | 0,425 | R_EC | 0,428 | R_EC | 0,371 | | R_EC | 0,271 | R_EC | 0,137 | R_EC | 0,214 |
| R_PreS | 0,500 | R_PreS | 0,547 | R_PreS | 0,429 | | R_PreS | 0,200 | R_PreS | 0,214 | R_PreS | 0,093 |
| R_H | 0,000 | R_H | 0,000 | R_H | 0,411 | | R_H | 0,000 | R_H | 0,000 | R_H | 0,261 |
| R_ProS | 0,527 | R_ProS | 0,681 | R_ProS | 0,481 | | R_ProS | 0,323 | R_ProS | 0,431 | R_ProS | 0,227 |
| R_PeEc | 0,581 | R_PeEc | 0,404 | R_PeEc | 0,454 | | R_PeEc | 0,334 | R_PeEc | 0,044 | R_PeEc | 0,189 |
| R_STGa | 0,475 | R_STGa | 0,619 | R_STGa | 0,437 | | R_STGa | 0,257 | R_STGa | 0,181 | R_STGa | 0,224 |
| R_PBelt | 0,569 | R_PBelt | 0,702 | R_PBelt | 0,556 | | R_PBelt | 0,319 | R_PBelt | 0,331 | R_PBelt | 0,308 |
| R_A5 | 0,535 | R_A5 | 0,670 | R_A5 | 0,547 | | R_A5 | 0,222 | R_A5 | 0,248 | R_A5 | 0,269 |
| R_PHA1 | 0,401 | R_PHA1 | 0,488 | R_PHA1 | 0,480 | | R_PHA1 | 0,210 | R_PHA1 | 0,148 | R_PHA1 | 0,142 |
| R_PHA3 | 0,408 | R_PHA3 | 0,436 | R_PHA3 | 0,494 | | R_PHA3 | 0,298 | R_PHA3 | 0,103 | R_PHA3 | 0,185 |
| R_STSda | 0,517 | R_STSda | 0,619 | R_STSda | 0,553 | | R_STSda | 0,240 | R_STSda | 0,196 | R_STSda | 0,183 |
| R_STSdp | 0,480 | R_STSdp | 0,659 | R_STSdp | 0,560 | | R_STSdp | 0,290 | R_STSdp | 0,290 | R_STSdp | 0,289 |
| R_STSvp | 0,444 | R_STSvp | 0,697 | R_STSvp | 0,502 | | R_STSvp | 0,262 | R_STSvp | 0,297 | R_STSvp | 0,204 |
| R_TGd | 0,555 | R_TGd | 0,527 | R_TGd | 0,376 | | R_TGd | 0,270 | R_TGd | 0,088 | R_TGd | 0,082 |
| R_TE1a | 0,406 | R_TE1a | 0,567 | R_TE1a | 0,447 | | R_TE1a | 0,177 | R_TE1a | 0,107 | R_TE1a | 0,123 |
| R_TE1p | 0,402 | R_TE1p | 0,593 | R_TE1p | 0,460 | | R_TE1p | 0,184 | R_TE1p | 0,243 | R_TE1p | 0,201 |
| R_TE2a | 0,558 | R_TE2a | 0,575 | R_TE2a | 0,421 | | R_TE2a | 0,247 | R_TE2a | 0,263 | R_TE2a | 0,103 |
| R_TF | 0,501382619 | R_TF | 0,481 | R_TF | 0,513 | | R_TF | 0,366 | R_TF | 0,067 | R_TF | 0,190 |
| R_TE2p | 0,518205303 | R_TE2p | 0,452 | R_TE2p | 0,468 | | R_TE2p | 0,319 | R_TE2p | 0,185 | R_TE2p | 0,075 |
| R_PHT | 0,462 | R_PHT | 0,572 | R_PHT | 0,510 | | R_PHT | 0,268 | R_PHT | 0,306 | R_PHT | 0,177 |
| R_PH | 0,429 | R_PH | 0,589 | R_PH | 0,460 | | R_PH | 0,270 | R_PH | 0,277 | R_PH | 0,177 |
| R_TPOJ1 | 0,518 | R_TPOJ1 | 0,675 | R_TPOJ1 | 0,498 | | R_TPOJ1 | 0,220 | R_TPOJ1 | 0,256 | R_TPOJ1 | 0,247 |
| R_TPOJ2 | 0,508 | R_TPOJ2 | 0,652 | R_TPOJ2 | 0,455 | | R_TPOJ2 | 0,229 | R_TPOJ2 | 0,341 | R_TPOJ2 | 0,257 |
| R_TPOJ3 | 0,481 | R_TPOJ3 | 0,671 | R_TPOJ3 | 0,479 | | R_TPOJ3 | 0,230 | R_TPOJ3 | 0,367 | R_TPOJ3 | 0,252 |
| R_DVT | 0,519 | R_DVT | 0,653 | R_DVT | 0,468 | | R_DVT | 0,274 | R_DVT | 0,340 | R_DVT | 0,221 |
| R_PGp | 0,381 | R_PGp | 0,557 | R_PGp | 0,436 | | R_PGp | 0,312 | R_PGp | 0,348 | R_PGp | 0,217 |
| R_IP2 | 0,559 | R_IP2 | 0,703 | R_IP2 | 0,485 | | R_IP2 | 0,349 | R_IP2 | 0,393 | R_IP2 | 0,242 |

| | | | | | | | | | | | | |
|---|---|---|---|---|---|---|---|---|---|---|---|---|
| R_IP1 | 0,455 | R_IP1 | 0,628 | R_IP1 | 0,470 | | R_IP1 | 0,415 | R_IP1 | 0,360 | R_IP1 | 0,279 |
| R_IP0 | 0,436 | R_IP0 | 0,640 | R_IP0 | 0,487 | | R_IP0 | 0,301 | R_IP0 | 0,303 | R_IP0 | 0,231 |
| R_PFop | 0,487 | R_PFop | 0,679 | R_PFop | 0,543 | | R_PFop | 0,356 | R_PFop | 0,303 | R_PFop | 0,268 |
| R_PF | 0,557 | R_PF | 0,632 | R_PF | 0,526 | | R_PF | 0,367 | R_PF | 0,405 | R_PF | 0,368 |
| R_PFm | 0,469 | R_PFm | 0,648 | R_PFm | 0,499 | | R_PFm | 0,459 | R_PFm | 0,484 | R_PFm | 0,321 |
| R_PGi | 0,429 | R_PGi | 0,664 | R_PGi | 0,464 | | R_PGi | 0,367 | R_PGi | 0,464 | R_PGi | 0,214 |
| R_PGs | 0,423 | R_PGs | 0,660 | R_PGs | 0,445 | | R_PGs | 0,456 | R_PGs | 0,461 | R_PGs | 0,246 |
| R_V6A | 0,488 | R_V6A | 0,633 | R_V6A | 0,474 | | R_V6A | 0,394 | R_V6A | 0,317 | R_V6A | 0,243 |
| R_VMV1 | 0,461 | R_VMV1 | 0,659 | R_VMV1 | 0,472 | | R_VMV1 | 0,354 | R_VMV1 | 0,388 | R_VMV1 | 0,270 |
| R_VMV3 | 0,468 | R_VMV3 | 0,627 | R_VMV3 | 0,490 | | R_VMV3 | 0,278 | R_VMV3 | 0,354 | R_VMV3 | 0,307 |
| R_PHA2 | 0,451 | R_PHA2 | 0,477 | R_PHA2 | 0,494 | | R_PHA2 | 0,318 | R_PHA2 | 0,198 | R_PHA2 | 0,198 |
| R_V4t | 0,507 | R_V4t | 0,661 | R_V4t | 0,428 | | R_V4t | 0,271 | R_V4t | 0,343 | R_V4t | 0,273 |
| R_FST | 0,481 | R_FST | 0,680 | R_FST | 0,489 | | R_FST | 0,237 | R_FST | 0,353 | R_FST | 0,200 |
| R_V3CD | 0,492 | R_V3CD | 0,639 | R_V3CD | 0,505 | | R_V3CD | 0,470 | R_V3CD | 0,479 | R_V3CD | 0,343 |
| R_LO3 | 0,448 | R_LO3 | 0,629 | R_LO3 | 0,440 | | R_LO3 | 0,412 | R_LO3 | 0,417 | R_LO3 | 0,287 |
| R_VMV2 | 0,439 | R_VMV2 | 0,644 | R_VMV2 | 0,491 | | R_VMV2 | 0,286 | R_VMV2 | 0,308 | R_VMV2 | 0,246 |
| R_31pd | 0,510 | R_31pd | 0,692 | R_31pd | 0,569 | | R_31pd | 0,348 | R_31pd | 0,402 | R_31pd | 0,287 |
| R_31a | 0,371 | R_31a | 0,635 | R_31a | 0,547 | | R_31a | 0,247 | R_31a | 0,282 | R_31a | 0,301 |
| R_VVC | 0,453 | R_VVC | 0,582 | R_VVC | 0,473 | | R_VVC | 0,304 | R_VVC | 0,294 | R_VVC | 0,243 |
| R_25 | 0,485 | R_25 | 0,348 | R_25 | 0,404 | | R_25 | 0,390 | R_25 | 0,187 | R_25 | 0,147 |
| R_s32 | 0,462 | R_s32 | 0,474 | R_s32 | 0,515 | | R_s32 | 0,373 | R_s32 | 0,195 | R_s32 | 0,197 |
| R_pOFC | 0,513 | R_pOFC | 0,306 | R_pOFC | 0,446 | | R_pOFC | 0,297 | R_pOFC | 0,093 | R_pOFC | 0,132 |
| R_PoI1 | 0,438 | R_PoI1 | 0,565 | R_PoI1 | 0,516 | | R_PoI1 | 0,149 | R_PoI1 | 0,168 | R_PoI1 | 0,176 |
| R_Ig | 0,493 | R_Ig | 0,643 | R_Ig | 0,504 | | R_Ig | 0,265 | R_Ig | 0,418 | R_Ig | 0,330 |
| R_FOP5 | 0,472 | R_FOP5 | 0,687 | R_FOP5 | 0,591 | | R_FOP5 | 0,317 | R_FOP5 | 0,257 | R_FOP5 | 0,269 |
| R_p10p | 0,392 | R_p10p | 0,595 | R_p10p | 0,393 | | R_p10p | 0,201 | R_p10p | 0,302 | R_p10p | 0,256 |
| R_p47r | 0,474 | R_p47r | 0,629 | R_p47r | 0,469 | | R_p47r | 0,243 | R_p47r | 0,379 | R_p47r | 0,185 |
| R_TGv | 0,604 | R_TGv | 0,458 | R_TGv | 0,515 | | R_TGv | 0,237 | R_TGv | 0,108 | R_TGv | 0,247 |
| R_MBelt | 0,500 | R_MBelt | 0,648 | R_MBelt | 0,508 | | R_MBelt | 0,302 | R_MBelt | 0,344 | R_MBelt | 0,236 |
| R_LBelt | 0,569 | R_LBelt | 0,696 | R_LBelt | 0,533 | | R_LBelt | 0,326 | R_LBelt | 0,337 | R_LBelt | 0,323 |

| | | | | | | | | | | | | |
|---|---|---|---|---|---|---|---|---|---|---|---|---|
| R_A4 | 0,504 | R_A4 | 0,675 | R_A4 | 0,565 | | R_A4 | 0,193 | R_A4 | 0,297 | R_A4 | 0,255 |
| R_STSva | 0,529 | R_STSva | 0,656 | R_STSva | 0,491 | | R_STSva | 0,203 | R_STSva | 0,266 | R_STSva | 0,201 |
| R_TE1m | 0,489 | R_TE1m | 0,622 | R_TE1m | 0,452 | | R_TE1m | 0,364 | R_TE1m | 0,210 | R_TE1m | 0,208 |
| R_PI | 0,388 | R_PI | 0,457 | R_PI | 0,184 | | R_PI | 0,204 | R_PI | 0,193 | R_PI | 0,108 |
| R_a32pr | 0,391 | R_a32pr | 0,615 | R_a32pr | 0,587 | | R_a32pr | 0,340 | R_a32pr | 0,299 | R_a32pr | 0,211 |
| R_p24 | 0,394 | R_p24 | 0,544 | R_p24 | 0,522 | | R_p24 | 0,258 | R_p24 | 0,162 | R_p24 | 0,314 |
| L_accumbens | 0,524 | L_accumbens | 0,408 | L_accumbens.area | 0,436 | | L_accumbens | 0,449 | L_accumbens | 0,168 | L_accumbens.area | 0,117 |
| L_amygdala | 0,485 | L_amygdala | 0,568 | L_amygdala | 0,352 | | L_amygdala | 0,314 | L_amygdala | 0,244 | L_amygdala | 0,084 |
| L_caudate | 0,385 | L_caudate | 0,603 | L_caudate | 0,543 | | L_caudate | 0,178 | L_caudate | 0,156 | L_caudate | 0,116 |
| L_hippocampus | 0,427 | L_hippocampus | 0,496 | L_hippocampus | 0,371 | | L_hippocampus | 0,254 | L_hippocampus | 0,068 | L_hippocampus | 0,100 |
| L_pallidum | 0,354 | L_pallidum | 0,505 | L_pallidum | 0,480 | | L_pallidum | 0,352 | L_pallidum | 0,263 | L_pallidum | 0,165 |
| L_putamen | 0,402 | L_putamen | 0,616 | L_putamen | 0,551 | | L_putamen | 0,171 | L_putamen | 0,196 | L_putamen | 0,202 |
| L_thalamus | 0,342 | L_thalamus | 0,511 | L_thalamus | 0,486 | | L_thalamus | 0,101 | L_thalamus | 0,157 | L_thalamus_proper | 0,107 |
| L_ventraldc | 0,344 | L_ventraldc | 0,520 | L_ventraldc | 0,429 | | L_ventraldc | 0,173 | L_ventraldc | 0,149 | L_ventraldc | 0,069 |
| R_accumbens | 0,553 | R_accumbens | 0,387 | R_accumbens | 0,449 | | R_accumbens | 0,388 | R_accumbens | 0,101 | R_accumbens.area | 0,173 |
| R_amygdala | 0,479 | R_amygdala | 0,556 | R_amygdala | 0,407 | | R_amygdala | 0,393 | R_amygdala | 0,180 | R_amygdala | 0,095 |
| R_caudate | 0,491 | R_caudate | 0,591 | R_caudate | 0,544 | | R_caudate | 0,272 | R_caudate | 0,201 | R_caudate | 0,105 |
| R_hippocampus | 0,416 | R_hippocampus | 0,525 | R_hippocampus | 0,344 | | R_hippocampus | 0,167 | R_hippocampus | 0,099 | R_hippocampus | 0,153 |
| R_pallidum | 0,300 | R_pallidum | 0,463 | R_pallidum | 0,438 | | R_pallidum | 0,201 | R_pallidum | 0,190 | R_pallidum | 0,136 |
| R_putamen | 0,444 | R_putamen | 0,616 | R_putamen | 0,541 | | R_putamen | 0,170 | R_putamen | 0,051 | R_putamen | 0,180 |
| R_thalamus | 0,364 | R_thalamus | 0,509 | R_thalamus | 0,482 | | R_thalamus | 0,128 | R_thalamus | 0,135 | R_thalamus_proper | 0,138 |
| R_ventraldc | 0,386 | R_ventraldc | 0,504 | R_ventraldc | 0,449 | | R_ventraldc | 0,126 | R_ventraldc | 0,112 | R_ventraldc | 0,076 |

**Supplementary Table S6** Results of multiple regression analyses for nodal efficiency. The areas with non-zero effect sizes identified via elastic-net in one sample (left column) have been used as predictors for general intelligence in all other samples. Depicted are the coefficients of determination $R^2$ and the respective p-value (uncorrected for multiple comparisons). Control variables were age, sex, age*sex, age², age²*sex, and handedness. Predictors refers to the number of predictors identified in the sample.

| | predictors | HCP day 1 | | HCP day 2 | | HCP day 1 ses. 1 | | HCP day 1 ses. 2 | | HCP day 1&2 | | NKI (TR=645) | | NKI (TR=1400) | | NKI (TR=2500) | | UMN | | RUB | |
|---|---|---|---|---|---|---|---|---|---|---|---|---|---|---|---|---|---|---|---|---|---|
| | | $R^2$ | *p* | $R^2$ | *p* | $R^2$ | *p* | $R^2$ | *p* | $R^2$ | *p* | $R^2$ | *p* | $R^2$ | *p* | $R^2$ | *p* | $R^2$ | *p* | $R^2$ | *p* |
| **HCP day 1** | 91 | .29 | <.001 | .15 | <.001 | .19 | <.001 | .20 | <.001 | .16 | <.001 | .29 | .038 | .33 | .028 | .26 | .209 | .41 | .027 | .17 | .301 |
| **HCP day 2** | 33 | .08 | <.001 | .15 | <.001 | .08 | <.001 | .09 | <.001 | .06 | .002 | .16 | .002 | .13 | .054 | .11 | .112 | .15 | .147 | .06 | .501 |
| **HCP day 1 ses. 1** | 56 | .14 | <.001 | .10 | <.001 | .23 | <.001 | .14 | <.001 | .07 | .046 | .20 | .025 | .27 | <.001 | .16 | .276 | .23 | .209 | .09 | .606 |
| **HCP day 1 ses. 2** | 53 | .14 | <.001 | .10 | <.001 | .14 | <.001 | .23 | <.001 | .08 | .007 | .19 | .018 | .24 | .001 | .15 | .252 | .24 | .079 | .14 | .019 |
| **HCP day 1&2** | 40 | .09 | <.001 | .08 | <.001 | .10 | <.001 | .07 | <.001 | .17 | <.001 | .15 | .022 | .27 | <.001 | .14 | .077 | .16 | .302 | .10 | .034 |
| **NKI (TR = 645)** | 28 | .06 | <.001 | .03 | .155 | .05 | <.001 | .06 | <.001 | .03 | .277 | .28 | <.001 | .18 | <.001 | .15 | <.001 | .12 | .253 | .07 | .083 |
| **NKI (TR = 1400)** | 31 | .06 | .001 | .05 | .008 | .05 | .005 | .07 | <.001 | .04 | .250 | .16 | <.001 | .38 | <.001 | .15 | .002 | .23 | <.001 | .05 | .578 |
| **NKI (TR = 2500)** | 44 | .05 | .135 | .05 | .393 | .08 | <.001 | .08 | <.001 | .05 | .152 | .22 | <.001 | .28 | <.001 | .40 | <.001 | .12 | .886 | .06 | .863 |
| **UMN** | 31 | .06 | <.001 | .04 | .093 | .06 | <.001 | .10 | <.001 | .04 | .136 | .11 | .105 | .18 | <.001 | .07 | .658 | .40 | <.001 | .05 | .707 |
| **RUB** | 0 | - | - | - | - | - | - | - | - | - | - | - | - | - | - | - | - | - | - | - | - |

**Supplementary Table S7** Results of multiple regression analyses for nodal efficiency. The areas with non-zero effect sizes identified via elastic-net in one sample (left column) have been used as predictors for general intelligence in all other samples. Depicted is the number of significant predictors in the regression analyses along with its percentage compared to the number of predictors identified via elastic-net (uncorrected for multiple comparisons). Control variables were age, sex, age*sex, age², age²*sex, and handedness. Predictors refers to the number of predictors identified in the sample.

| | predictors | HCP day 1 | HCP day 2 | HCP day 1 ses. 1 | HCP day 1 ses. 2 | HCP day 1&2 | NKI (TR=645) | NKI (TR=1400) | NKI (TR=2500) | UMN | RUB |
|---|---|---|---|---|---|---|---|---|---|---|---|
| **HCP day 1** | 91 | 28 (31%) | 11 (12%) | 17 (19%) | 16 (18%) | 16 (18%) | 7 (8%) | 2 (2%) | 6 (7%) | 6 (7%) | 6 (7%) |
| **HCP day 2** | 33 | 3 (9%) | 15 (45%) | 4 (12%) | 4 (12%) | 5 (15%) | 2 (6%) | 3 (9%) | 3 (9%) | 1 (3%) | 1 (3%) |
| **HCP day 1 ses. 1** | 56 | 10 (18%) | 6 (11%) | 13 (23%) | 8 (14%) | 5 (9%) | 4 (7%) | 9 (16%) | 1 (2%) | 3 (5%) | 2 (4%) |
| **HCP day 1 ses. 2** | 53 | 16 (30%) | 8 (15%) | 10 (19%) | 18 (34%) | 6 (11%) | 5 (9%) | 5 (9%) | 1 (2%) | 3 (6%) | 5 (9%) |
| **HCP day 1&2** | 40 | 9 (23%) | 7 (18%) | 9 (23%) | 6 (15%) | 12 (30%) | 3 (8%) | 6 (15%) | 2 (5%) | 2 (5%) | 5 (13%) |
| **NKI (TR = 645)** | 28 | 5 (18%) | 3 (11%) | 4 (14%) | 5 (18%) | 5 (18%) | 6 (21%) | 1 (4%) | 1 (4%) | 3 (11%) | 2 (7%) |
| **NKI (TR = 1400)** | 31 | 3 (10%) | 3 (10%) | 1 (3%) | 3 (10%) | 1 (3%) | 2 (6%) | 11 (35%) | 3 (10%) | 6 (19%) | 2 (6%) |
| **NKI (TR = 2500)** | 44 | 2 (5%) | 2 (5%) | 4 (9%) | 4 (9%) | 3 (7%) | 6 (14%) | 7 (16%) | 15 (34%) | 1 (2%) | 2 (5%) |
| **UMN** | 31 | 3 (10%) | 3 (10%) | 2 (6%) | 6 (19%) | 3 (10%) | 3 (10%) | 5 (16%) | 1 (3%) | 11 (35%) | 2 (6%) |
| **RUB** | 0 | 0 | 0 | 0 | 0 | 0 | 0 | 0 | 0 | 0 | 0 |

**Supplementary Table S8** Results of multiple regression analyses for local clustering. The areas with non-zero effect sizes identified via elastic-net in one sample (left column) have been used as predictors for general intelligence in all other samples. Depicted are the coefficients of determination $R^2$ and the respective p-value (uncorrected for multiple comparisons). Control variables were age, sex, age*sex, age², age²*sex, and handedness.Predictors refers to the number of predictors identified in the sample.

| | predictors | HCP day 1 | | HCP day 2 | | HCP day 1 ses. 1 | | HCP day 1 ses. 2 | | HCP day 1&2 | | NKI (TR=645) | | NKI (TR=1400) | | NKI (TR=2500) | | UMN | | RUB | |
|---|---|---|---|---|---|---|---|---|---|---|---|---|---|---|---|---|---|---|---|---|---|
| | | $R^2$ | *p* | $R^2$ | *p* | $R^2$ | *p* | $R^2$ | *p* | $R^2$ | *p* | $R^2$ | *p* | $R^2$ | *p* | $R^2$ | *p* | $R^2$ | *p* | $R^2$ | *p* |
| **HCP day 1** | 52 | .21 | <.001 | .08 | .002 | .11 | <.001 | .08 | .001 | .08 | .006 | .18 | .032 | .22 | .008 | .19 | .016 | .22 | .202 | .11 | .188 |
| **HCP day 2** | 48 | .08 | .001 | .20 | <.001 | .10 | <.001 | .07 | .007 | .06 | .160 | .17 | .046 | .19 | .022 | .19 | .007 | .18 | .390 | .08 | .522 |
| **HCP day 1 ses. 1** | 41 | .07 | <.001 | .06 | .037 | .20 | <.001 | .06 | .013 | .06 | .061 | .18 | .002 | .18 | .011 | .12 | .257 | .12 | .812 | .08 | .315 |
| **HCP day 1 ses. 2** | 17 | .04 | <.001 | .03 | .007 | .05 | <.001 | .09 | <.001 | .02 | .226 | .08 | .011 | .08 | .054 | .05 | .311 | .03 | .957 | .05 | .061 |
| **HCP day 1&2** | 50 | .08 | <.001 | .08 | .006 | .07 | .005 | .07 | .013 | .21 | <.001 | .21 | .002 | .20 | .019 | .18 | .022 | .17 | .637 | .11 | .181 |
| **NKI (TR = 645)** | 27 | .05 | .002 | .04 | .022 | .06 | <.001 | .05 | .005 | .05 | .003 | .27 | <.001 | .17 | <.001 | .09 | .115 | .12 | .232 | .07 | .102 |
| **NKI (TR = 1400)** | 39 | .04 | .350 | .06 | .013 | .06 | .019 | .05 | .028 | .04 | .282 | .16 | .008 | .37 | <.001 | .15 | .015 | .17 | .162 | .08 | .292 |
| **NKI (TR = 2500)** | 16 | .03 | .018 | .02 | .210 | .02 | .108 | .03 | .032 | .03 | .030 | .07 | .037 | .13 | <.001 | .22 | <.001 | .05 | .631 | .04 | .111 |
| **UMN** | 2 | .01 | .072 | .00 | .162 | .00 | .225 | .01 | .020 | .00 | .540 | .03 | .005 | .01 | .114 | .00 | .793 | .07 | <.001 | .00 | .760 |
| **RUB** | 0 | - | - | - | - | - | - | - | - | - | - | - | - | - | - | - | - | - | - | - | - |

**Supplementary Table S9** Results of multiple regression analyses for local clustering. The areas with non-zero effect sizes identified via elastic-net in one sample (left column) have been used as predictors for general intelligence in all other samples. Depicted is the number of significant predictors in the regression analyses along with its percentage compared to the number of predictors identified via elastic-net (uncorrected for multiple comparisons). Control variables were age, sex, age*sex, age², age²*sex, and handedness.Predictors refers to the number of predictors identified in the sample.

| | predictors | | | | | | | | | | |
|---|---|---|---|---|---|---|---|---|---|---|---|
| | | **HCP day 1** | **HCP day 2** | **HCP day 1 ses. 1** | **HCP day 1 ses. 2** | **HCP day 1&2** | **NKI (TR=645)** | **NKI (TR=1400)** | **NKI (TR=2500)** | **UMN** | **RUB** |
| **HCP day 1** | 52 | 21 (40%) | 4 (8%) | 6 (12%) | 5 (10%) | 5 (10%) | 3 (6%) | 5 (10%) | 3 (6%) | 3 (6%) | 5 (10%) |
| **HCP day 2** | 48 | 4 (8%) | 21 (44%) | 8 (17%) | 3 (6%) | 4 (8%) | 3 (6%) | 4 (8%) | 3 (6%) | 3 (6%) | 2 (4%) |
| **HCP day 1 ses. 1** | 41 | 3 (7%) | 3 (7%) | 20 (49%) | 2 (5%) | 4 (10%) | 5 (12%) | 4 (10%) | 3 (7%) | 1 (2%) | 2 (5%) |
| **HCP day 1 ses. 2** | 17 | 2 (12%) | 2 (12%) | 2 (12%) | 6 (35%) | 2 (12%) | 1 (6%) | 2 (12%) | 1 (6%) | 1 (6%) | 2 (12%) |
| **HCP day 1&2** | 50 | 6 (12%) | 6 (12%) | 8 (16%) | 3 (6%) | 30 (60%) | 8 (16%) | 2 (4%) | 3 (6%) | 2 (4%) | 3 (6%) |
| **NKI (TR = 645)** | 27 | 5 (19%) | 1 (4%) | 3 (11%) | 5 (19%) | 1 (4%) | 5 (19%) | 3 (11%) | 2(7%) | 2 (7%) | 2 (7%) |
| **NKI (TR = 1400)** | 39 | 0 (0%) | 3 (8%) | 1 (3%) | 4 (10%) | 3 (8%) | 1 (3%) | 11 (28%) | 3 (8%) | 3 (8%) | 4 (10%) |
| **NKI (TR = 2500)** | 16 | 2 (13%) | 1 (6%) | 1 (6%) | 2 (13%) | 2 (13%) | 2 (13%) | 4 (25%) | 9 (56%) | 0 (0%) | 0 (0%) |
| **UMN** | 2 | 1 (50%) | 0 (0%) | 0 (0%) | 1 (50%) | 0 (0%) | 1 (50%) | 1 (50%) | 0 (0%) | 2 (100%) | 0 (0%) |
| **RUB** | 0 | 0 | 0 | 0 | 0 | 0 | 0 | 0 | 0 | 0 | 0 |
